# Drift on holey landscapes as a dominant evolutionary process

**DOI:** 10.1101/2021.10.22.465488

**Authors:** Ned A. Dochtermann, Brady Klock, Derek A. Roff, Raphaël Royauté

## Abstract

An organism’s phenotype has been shaped by evolution but the specific processes have to be indirectly inferred for most species. For example, correlations among traits imply the historical action of correlated selection and, more generally, the expression and distribution of traits is expected to be reflective of the adaptive landscapes that have shaped a population. However, our expectations about how quantitative traits—like most behaviors, physiological processes, and life-history traits—should be distributed under different evolutionary processes is not clear. Here we show that genetic variation in quantitative traits is not distributed as would be expected under dominant evolutionary models. Instead, we found that genetic variation in quantitative traits across 6 phyla and 60 species (including both Plantae and Animalia) is consistent with evolution across high dimensional “holey landscapes”. This suggests that the leading conceptualizations and modeling of the evolution of trait integration fail to capture how phenotypes are shaped and that traits are integrated in a manner contrary to predictions of dominant evolutionary theory. Our results demonstrate that our understanding of how evolution has shaped phenotypes remains incomplete and these results provide a starting point for reassessing the relevance of existing evolutionary models.

**Significance Statement:** We found that empirical estimations of how quantitative genetic variation is distributed do not correspond to typical Gaussian representations of fitness landscapes. These Gaussian landscapes underpin major areas of evolutionary biology and how selection is estimated in natural populations. Rather than being consistent with evolution on Gaussian landscapes, empirical estimates of genetic variation are, instead, consistent with evolution on high-dimensional “holey” landscapes. These landscapes represent situations where specific combinations of trait values are either viable or not and populations randomly drift among the viable combinations. This finding suggests that we have substantially misunderstood how selection actually shapes populations and thus how evolution typically proceeds.

## Introduction

Our understanding of selection has been strongly shaped by Sewall Wright’s conceptualization of an evolutionary landscape, with populations moving from areas of low fitness to areas of higher fitness (1, 2). While the one and two trait landscapes Wright originally described have been criticized as unrealistic, including by Wright himself (1), the general metaphor has nonetheless guided much of evolutionary thought (3).

For quantitative traits, like many aspects of physiology, behavior, and morphology, Wright’s metaphor has been extended to complex topographies with ridges or tunnels of high fitness (4–6). Mathematical implementations of these landscapes have led to insights into how topography shapes the distribution of genetic (co)variance (7–9). Gaussian landscapes influenced by Wright’s metaphor also underpin the statistical approaches that are used to estimate the action of stabilizing selection in natural populations (10).

How the topography of evolutionary landscapes affects sequence evolution at the genomic level has garnered similar interest (11). While research into quantitative traits has focused on relatively simple landscapes (e.g. 7, 8, 9, 12-14), research regarding sequence evolution spans simple single peak Gaussian “Fujiyama landscapes”, to “badlands landscapes” (Fig 1A & 1B (15)), to abstract high-dimensional “holey landscapes” (Fig 1C (16)). An important conclusion from this research is that evolutionary dynamics on simple landscapes often fail to properly predict evolution on higher dimensional landscapes. Empirical research into quantitative traits has been slow to incorporate this need for a higher-dimensional perspective.

**Figure 1.**
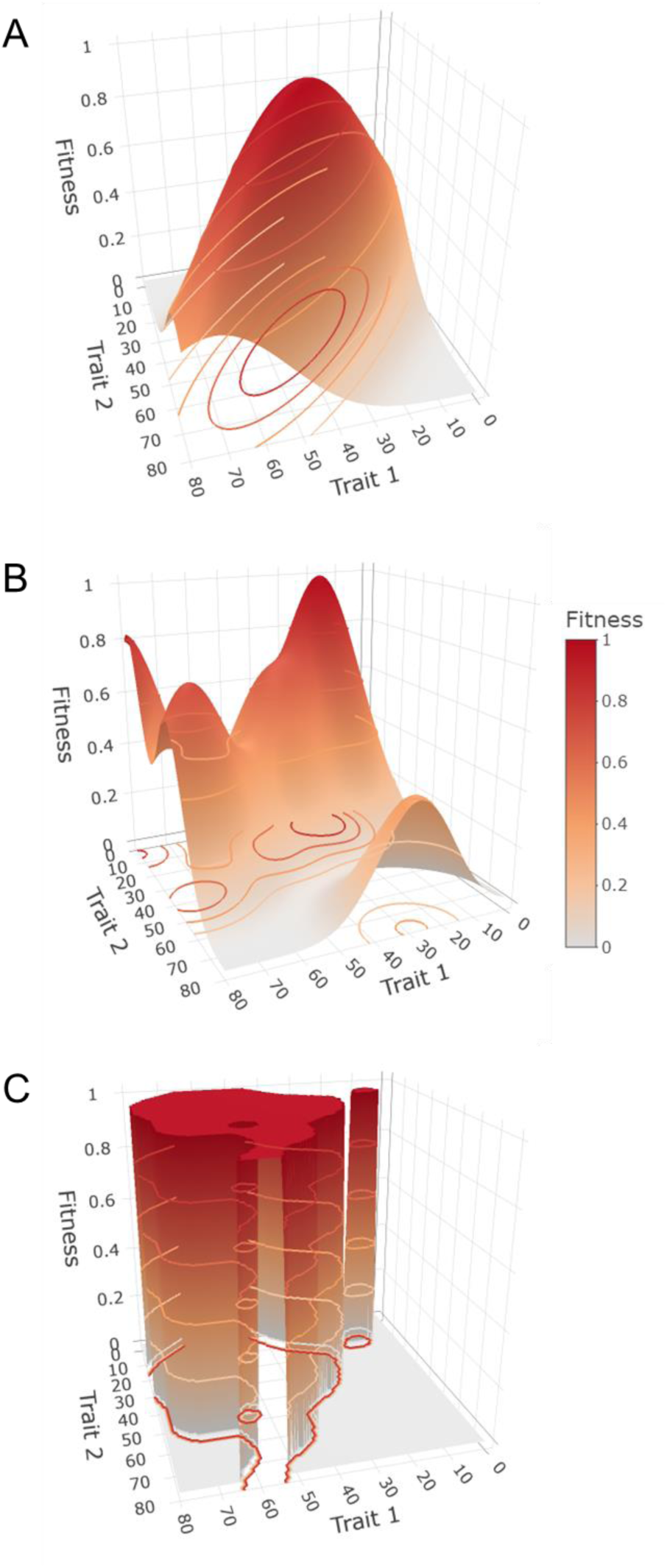
Example fitness landscapes. Redder colors correspond to higher fitness. A. A simple Gaussian, single peak Fujiyama landscape with a single optimum. B. A more rugged landscape with multiple local optima and a single global optimum. C. A simplified Holey landscape where combinations of values correspond to high, average, fitness (1) or low (0) fitness. This landscape has a large cluster of viable phenotypes with a hole and irregular borders and a smaller cluster of viable phenotypes at low trait values.

Perhaps most conceptually unfamiliar and unintuitive to researchers focused on quantitative traits are holey landscapes (Fig 1C (16)). Holey landscapes are high-dimensional evolutionary landscapes that consist of trait combinations that are either of average fitness or that are inviable (16, 17). This results in flat landscapes with holes at inviable or low fitness phenotypes (Fig 1C). This topography stems from the multivariate nature of phenotypes: while there may be continuous fitness differences in two dimensions, fitness gradients will create holes in the landscape and peaks will average out when additional traits are considered. Unfortunately, predictions about quantitative trait evolution on holey landscapes are not clear and have rarely been pursued, (e.g. 18).

Even more importantly, it is unknown what the topography of landscapes is for natural populations. While portions of selection surfaces and landscapes can be directly estimated (10, 19), these estimates may differ from the full landscape due to several factors. These include: the omission of fitness affecting traits (20), incomplete estimation of fitness (21, 22), and insufficient power to estimate non-linear selection coefficients (23).

An alternative to direct estimation of landscape topography is to infer landscape topography from observed trait (co)variances. For example, low additive genetic variation is suggestive of stabilizing or directional selection (24) and additive genetic correlations are expected to emerge from correlational selection (e.g. 4, 5). An ability to gain an understanding of the topography of landscapes based on observed variation would greatly further our understanding of how selection is realized in natural populations.

Here we used a simulation model to examine how evolution on different landscapes shapes genetic (co)variation. We modeled populations as evolving solely via drift, evolving via adaptation on Gaussian fitness landscapes stemming from Wright’s metaphor, or as evolving on holey landscapes. While many additional implementations of Wright’s metaphor exist (reviewed by 25), we focused on Gaussian landscapes with single peaks. We made this choice because this implementation of the metaphor connects to statistical approaches to estimating selection in natural populations (10, 26, 27) and to models of how genetic (co)variances evolve (7, 9, 28). These models allowed us to generate testable predictions for how the structure of additive genetic variances and covariances (**G**) are shaped by different landscape topographies. We then compared these modeled outcomes to 181 estimates of **G**, representing 60 species from 6 phyla, including both plants and animals. We show that the observed distribution of genetic variation is ultimately more consistent with evolution on holey landscapes than on Wrightian landscapes, indicating our models of phenotypic evolution should be re-evaluated.

## Results

### Model outcomes

When evolving on holey landscapes, simulated populations lost greater relative variation in the non-dominant dimensions as compared to when evolving on simple Gaussian landscapes or when subject solely to drift (Fig 2; Fig S5). The ratio of genetic variation in the second versus first dimension (*λ*_2_⁄*λ*_1_) significantly differed depending on selection regime (F_4,1245_ = 368, p << 0.01; Fig 2). Populations experiencing either just drift or evolving on Gaussian landscapes maintained a more even amount of variation across dimensions compared to those evolving on holey landscapes (i.e. higher *λ*_2_⁄*λ*_1_ all post-hoc comparisons p < 0.001; Fig 2, Table S3). All populations evolving on holey landscapes exhibited similar *λ*_2_⁄*λ*_1_ ratios regardless of *p* (all post-hoc comparisons of outcomes for holey landscapes: p > 0.05; Fig 2, Table S3). This similarity is likely due to the observation elsewhere that, when *p* is greater than ½*^k^*, a cluster of viable phenotypic values—and therefore phenotypic space exists—through which a population can drift (18, 29). Given that *k* here was 10, this condition was satisfied.

**Figure 2.**
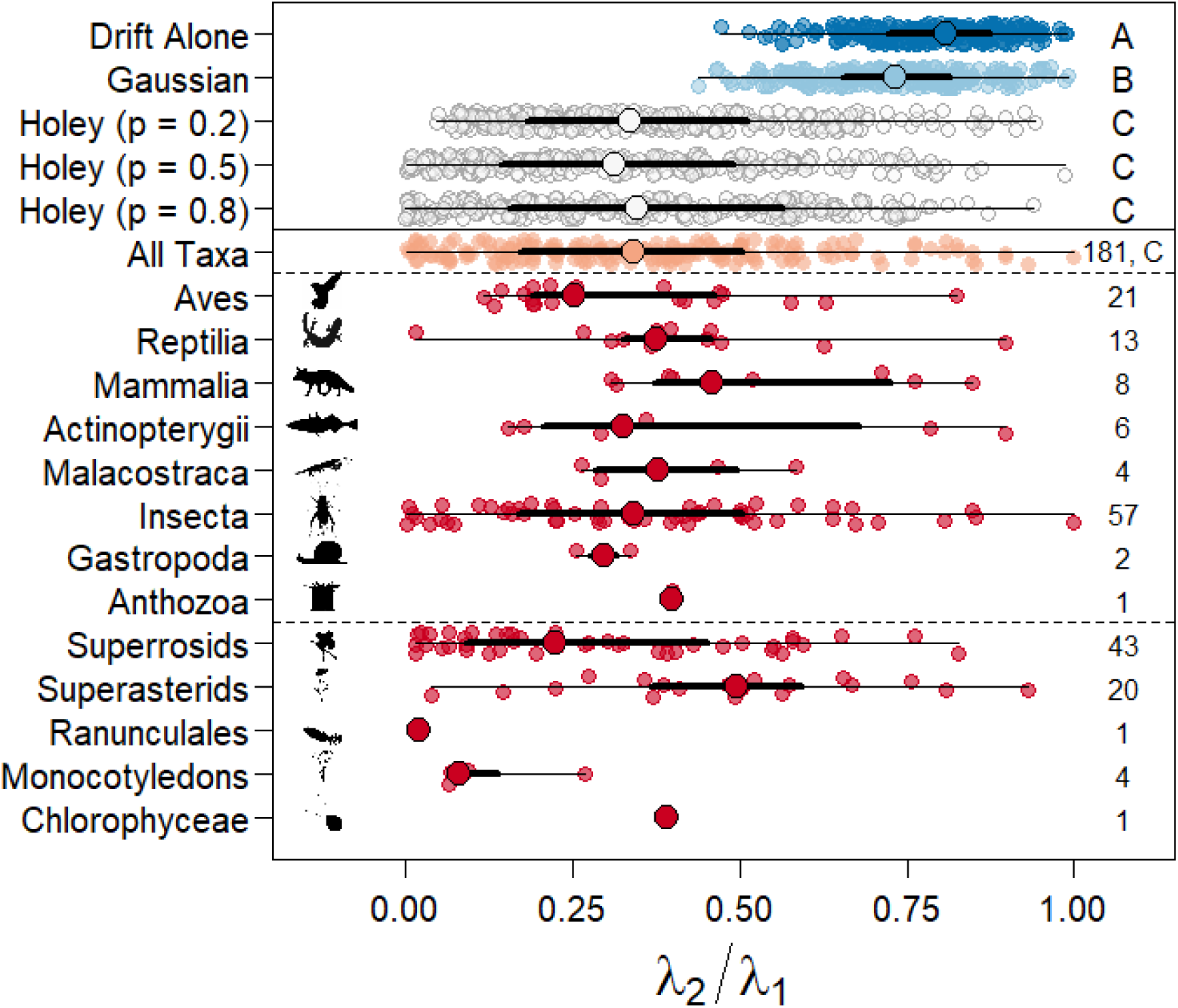
Modified “Orchard plot” of *λ*_2_⁄*λ*_1_ values for simulated (above solid line) and observed **G** matrices. *Trunks* (large points) are the medians for the specified group (e.g. Gaussian landscapes or Insecta), *branches* (thick lines) are interquartile ranges, *twigs* (thin lines) give the full range of values, and *fruits* (smaller points) are individual estimates within a simulation or taxonomic group. Rightmost letters correspond to statistical significance—or lack thereof—of comparisons of ratios among simulations. Datasets sharing letters did not significantly differ (Table S3). Populations evolving due to drift alone had a significantly higher ratio than observed for either stabilizing selection or evolution on any of the holey landscapes. Populations evolving on holey landscapes also had lower ratios than those experiencing stabilizing selection but did not differ from each other. Rightmost numbers are the number of estimates available via literature search. (organism silhouettes courtesy of phylopic.org, Public Domain Mark 1 licenses or CCA 3.0; Chlorophyceae: S.A. Muñoz-Gómez, Superrosid: D.J. Bruzzese, Superasterid: T.M. Keesey & Nadiatalent).

While a modest difference, populations evolving due to drift alone also exhibited a significantly greater ratio than populations evolving on Gaussian landscapes (difference = 0.06, p = 0.002; Fig 2, Table S3). This magnitude of a difference is unlikely to be biologically important or detectable in natural populations and instead is likely only detectable here due to the high power afforded by simulations. These differences were consistent across multiple approaches to summarizing **G** and are robust to conditions of the simulations (Tables S4 – S7, Figs S6 – S8). Interestingly, examination of single population outcomes suggests that the outcomes observed for populations evolving on Gaussian landscapes stem from the populations becoming trapped at local optima (e.g. Fig S4).

These modeling results produce the general prediction that greater relative variation in multiple dimensions is maintained when populations evolve on Gaussian landscapes than when evolving on holey landscapes. Put another way, evolving on holey landscapes is predicted to result in a large decrease in variation from the dominant to subsequent dimensions and, consequently, a lower *λ*_2_⁄*λ*_1_ value (Fig 2).

### Observed outcomes

Across all taxa, average *λ*_2_⁄*λ*_1_ was 0.36 (sd: 0.23, Fig 2). This estimate is consistent with and statistically indistinguishable from those observed for simulated populations evolving on holey landscapes (t_df:17.275_ = 0.32, 1.20, −0.05, p > 0.2 (all) versus holey landscapes with *p* = 0.2, 0.5, and 0.8 respectively; Fig 2, Table S10) and substantially less than observed for simulated populations that evolved on Gaussian landscapes or via drift alone (t_df:17.275_ = - 12.42, −14.55 respectively, p < 0.001 (both)).

While some individual estimates at the species level exhibited high *λ*_2_⁄*λ*_1_ values (Fig 2), phylogeny explained little variation in these values (phylogenetic heritability = 0.05; Table S9). As was the case across all taxa, median *λ*_2_⁄*λ*_1_ values for each taxonomic Class (or comparable level clade) were consistently lower than expected if evolution occurred on Gaussian landscapes or via drift alone (Fig 2). Instead, these results are strongly consistent with evolution on holey landscapes.

## Discussion

Wright’s conceptual model of adaptive landscapes has been key to our understanding of how selection acts. However, our results suggest something substantively different is occurring for many populations and species: the observed pattern of variation across taxa suggests that classic models of the evolution of quantitative traits are not the predominant force that has shaped the distribution of genetic variation. Instead, observed patterns of quantitative genetic variation are more consistent with drift across holey landscapes (Fig 2).

Our implementation of Wright’s metaphor represents only one of many possible evolutionary models (25). It is possible that unmodeled alternative landscapes may produce populations for which variation is distributed in a manner similar to holey landscapes and empirical estimates. As such, despite the strong similarity between empirical estimates of the structure of **G** and those from simulated populations that have evolved on holey landscapes, questions remain as to whether alternative landscapes are similarly likely. Importantly, and as mentioned previously, much of the exploration of evolution of quantitative traits has focused on simple landscapes like we have implemented here. Thus, it also is an open question what different models of selection “look” like when implemented for higher-dimensional phenotypes. For example, rugged landscapes of high-dimensionality may give rise to holey landscapes as peaks average out and valleys are inviable. Alternatively, but not mutually exclusively, Pareto-optimality may contribute to fitness planes (see below). Nonetheless, the close correspondence between empirical data and populations simulated as evolving on holey landscapes suggest that our understanding of quantitative trait evolution remains incomplete.

Much of the theoretical development of holey landscapes focused on the ability of populations to traverse genomic sequence differences via drift, with some sequences being inviable (e.g. due to missense differences in coding regions). How this extended to quantitative traits was less clear. Our simulation model provides one approach to applying the holey landscape framework to quantitative traits, treating each trait as a threshold character (30). Other approaches to modeling quantitative traits on holey landscapes and evolution in response to these versions, such as the generalized Russian roulette model (17), may produce different outcomes. Moreover, the topography of our modeled holey landscapes may also come about through other mechanisms. The flat surfaces of holey landscapes might instead emerge through similar processes as recent discussions of the Pareto optimization of traits (31, 32). Under Pareto optimization across just three traits a flat surface—the Pareto front—connects single trait × environment optima (i.e. “archetypes” (32)). Likewise, holes may come about not due to simple inviable trait combinations but, instead, rugged landscapes can create steep fitness declines and consequent holes in the overall landscape.

It is also important to recognize that the support for evolution on holey landscapes that we found does not preclude subsets of traits from having evolved on Gaussian landscapes. Indeed, stabilizing selection has been observed in natural populations (23), though understanding its general strength even on a case-by-case basis is confounded with methodological problems (33, 34). Regardless, our finding that observed patterns of quantitative genetic variation across taxonomic groups are not consistent with traditional evolutionary models stands.

This disconnect between observed patterns of multivariate variation and expectations under conventional models of selection suggests that Wright’s metaphor of landscapes—and the subsequent implementation of this metaphor as Gaussian surfaces— may have contributed to an incomplete understanding of how selection has shaped phenotypes. A potential contributor to this problem has been the lack of clear alternative explanations besides a simple null hypothesis of drift with no selection. Moving forward, clear development of additional alternative models of the action of selection and evolution in multivariate space are needed. This will allow the comparison of simulated populations to empirical data as we have done here.

Ultimately, our findings suggest that evolutionary biologists need to better consider the effects of high dimensionality as simple standard evolutionary models are not consistent with available data for quantitative data. Our findings also emphasize a need to consider *how* different landscapes might emerge. We are cautiously optimistic that novel development in high-throughput phenotyping (35, 36) will provide data that is better suited to answer the crucial questions we have raised.

## Materials & Methods

### Model Construction

We developed an individual variance components model (Methods, Fig S1 (37)) wherein individuals had phenotypes comprised of 10 traits (*k*), with each trait being highly heritable (h^2^ = 0.8), and initial genetic covariances between traits set at zero. Populations of individuals evolved on one of five landscapes: (i) a flat landscape where no selection occurred (i.e. drift alone), (ii) Gaussian landscapes where fitness for each pair of traits was characterized by a single peak but with correlational selection, and three (iii – v) implementations of holey landscapes differing by *p* (16, 17), the proportion of viable phenotypes in a holey landscape (*p* = 0.2, 0.5, and 0.8). Each of the modeling scenarios was simulated 250 times for populations of 7500 individuals and for 100 generations for each population. Full modeling details are provided in the Methods and all modeling code is available at https://github.com/DochtermannLab/Wright_vs_Holey.

### Model analysis

Following these simulations, the eigen structures of the resulting 1250 population genetic covariance matrices were compared. Because the simulated phenotypes consisted of 10 traits, it was the overall multivariate pattern of variation that was of interest rather than any specific single trait or pairwise combination (38). To do so, we calculated the ratio of each matrix’s second eigen value (λ_2_) to its dominant eigen value (i.e. *λ*_2_⁄*λ*_1_). The choice of using *λ*_2_⁄*λ*_1_was based on preliminary observations of modeling results. These results demonstrated that variation in populations that have evolved on holey landscapes was compressed into a primary dimension to a greater degree than observed for populations evolving under different conditions. *λ*_2_⁄*λ*_1_provides a better estimate of the compression of variance into a leading dimension than do other common metrics like the variation of the first eigen value to the sum of eigen values (i.e. *λ*_1_⁄∑ *λ*). For example, *λ*_1_⁄∑ *λ* could be low if the variation not captured by λ_1_ is equally distributed across all other dimensions, even if all other dimensions contained relatively little variation. The same scenario would produce a high value for *λ*_2_⁄*λ*_1_.

*λ*_2_⁄*λ*_1_ was then compared across the modeling scenarios using analysis of variance and Tukey post-hoc testing. Four alternative metrics for characterizing covariance matrices were consistent with the results for *λ*_2_⁄*λ*_1_ (see Supplementary Results). We also present the results of analyses of a broad range of starting conditions and model conditions in the Supplementary Results. These supplemental analyses covered much of the possible parameter space (Figures S12 – S14) and confirmed the robustness of the findings reported below.

### Observed patterns of genetic covariation

We next wanted to determine which of the modeled processes produced results consistent with observed patterns of trait integration. To do so, we conducted a literature review wherein we used Web of Science to search the journals American Naturalist, Ecology and Evolution, Evolution, Evolutionary Applications, Evolutionary Ecology, Genetics, Heredity, Journal of Evolutionary Biology, Journal of Heredity, Nature Ecology and Evolution, and the Proceedings of the Royal Society (B). We searched these journals using the terms “G matrix” on 14 May 2019, yielding a total of 272 articles. Each article was reviewed and estimated **G** matrices extracted if the article met inclusion criteria. For inclusion, an estimated **G** matrix must have been estimated for more than 2 traits (i.e. > 2 × 2), must have been reported as variances and covariances (i.e. not genetic correlations), and must not have been estimated for humans. Based on these inclusion criteria, we ended up with a dataset of 181 estimated **G** matrices from 60 articles (Fig S2). For each published **G** matrix, we estimated *λ*_2_⁄*λ*_1_. Study ID was included as a random effect to accommodate the estimation of different **G** matrices by group. This may not have fully captured the possibility of alternative fitness peaks by unknown grouping in underlying data.

As discussed above, the choice to use *λ*_2_⁄*λ*_1_ was an *a priori* decision based on examination of initial simulation runs. By focusing on a single predetermined metric we were able to conduct a confirmatory test of the concordance between published estimates of **G** and simulation outcomes. However, because *λ*_1_⁄∑ *λ* is more commonly reported, we also provide a graphical summary of simulation results and published estimates using this metric in the supplemental material (Fig S15). This summary, as with the primary results, demonstrate that published estimates of **G** more closely match populations evolving on holey landscapes than they do populations evolving on Gaussian landscapes.

## Acknowledgements

The authors thank B. de Bivort, A.J. Wilson, and J. Wright for helpful conversations. This work was supported by the US National Science Foundation grant 1557951 to N.A.D.

## Supplementary Materials

### Detailed Methods

#### Simulation Models

##### Model Construction

We developed an individual variance components model (Fig S1; sensu 1) wherein individuals had phenotypes comprised of 10 traits (*k*) and with each trait being highly heritable (h^2^ = 0.8) and initial genetic covariances between traits of 0. A high heritability was initially used to reduce the number of generations needed to determine the response of populations to selection. Genetic covariances were set to an initial value of zero to simulate a population under linkage equilibrium. Viability selection was applied based on fitness, which was determined either by location on a ten-dimensional holey landscape or on simple Gaussian landscapes with a single optimum per trait pair.

##### Holey Landscapes

For simulations evaluating holey landscapes, we simulated populations in which traits were inherited as though continuous but expressed categorically as one of two phenotypic variants (e.g. phenotype 0 versus 1 for trait 1). Specifically, at the start of simulations, we drew genotypes for each individual from a normal distribution with a mean of zero and standard deviation of 1. To these normally distributed genotypes, we added “environmental” values (µ = 0, all covariances = 0) to generate a phenotype with a heritability of 0.8. These continuously distributed phenotypic values were then transformed as one implementation of the holey landscape is based on the fitness of specific and discrete *combinations*. Specifically, the continuously distributed values were transformed to be a phenotype of 0 or 1, with a genotype < 0 being “0” and a genotype > 0 being “1” (Table S1).

The holey landscape for a specific simulation was then constructed by randomly assigning a fitness of 0 or 1 to the 1024 possible phenotypes (2*^k^*) trait combinations based on the parameter *p*. “*p*” was the probability that a trait combination had a fitness of 1 and corresponds to Gavrilets’ (2004) percolation parameter. We used three values of *p* in our simulation ranging from weak (*p* = 0.2), moderate (*p* = 0.5) and high (*p* =0.8). *p* can vary between 0 and 1, with values of 1 corresponding to a landscape where all trait combinations are viable and have a fitness of 1. As *p* approaches 0, few trait combinations are viable.

After the first generation, genotypes were drawn from a multivariate normal distribution based on the means and genetic variance-covariance matrix of the population that survived selection. Environmental contributions again had an average of 0 and no environmental correlation with a variance set to keep heritability at 0.8 (or other values during parameter exploration, below). The resulting phenotypic values were then converted to 0’s and 1’s as above. This approach to generating subsequent generations follows the structure of individual variance components models described by Roff (1). We used this individual variance components approach rather than an agent-based approach as the latter combined with the computational requirements of matching phenotypes to fitness under the holey landscape model was not amenable to simulation analysis.

**Table S1.** Example conversion of an underlying genotype to a phenotype under the two modelling scenarios. The same individual has a genotypic value for each of the 10 traits simulated (e.g. −0.918 for trait 10). To this, “environmental” contributions are added, taking heritability to 0.8. For Holey Landscape simulations, these phenotypic values are then converted to either 0 or 1 based on whether the phenotype is negative or positive.

|  | Trait |  |  |  |  |  |  |  |  |  |
| --- | --- | --- | --- | --- | --- | --- | --- | --- | --- | --- |
|  | 1 | 2 | 3 | 4 | 5 | 6 | 7 | 8 | 9 | 10 |
| Genotype | 0.008 | 0.770 | 0.477 | 0.112 | -0.512 | 0.751 | -1.752 | -0.944 | 0.030 | -0.918 |
| Environmental Contribution | 0.402 | -0.221 | 0.023 | 0.053 | 0.082 | -0.25 | 0.63 | 0.285 | -0.007 | 0.271 |
| Holey Landscape Phenotype | 1 | 1 | 1 | 1 | 0 | 1 | 0 | 0 | 1 | 0 |
| Gaussian Landscape Phenotype | 0.410 | 0.549 | 0.500 | 0.165 | -0.430 | 0.501 | -1.122 | -0.659 | 0.023 | -0.647 |

##### Gaussian (Wrightian) adaptive landscapes

For simulations evaluating Gaussian landscapes, we generated genotypes and phenotypes as above but without the categorical conversion (Table S1). We then generated random landscapes such that the optima (θ) for all traits was set to zero. The topography of the landscape for each pair of traits (e.g. ω_i,j_) was defined as

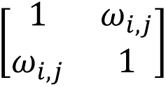 consistent with previous simulation studies examining the evolution of quantitative traits (reviewed by 2). This approach corresponds to single peak landscapes in any two dimensions. The forty-five ω*_i,j_* values that fully describe the landscape were generated using the LKJ onion method for constructing random correlation matrices with a pseudo-normal distribution of correlations where the average correlation is 0 (η = 1; Lewandowski et al. 2009; Fig S2). Using the LKJ onion method ensures that the full description of the landscape (ω) is positive semi-definite with feasible partial correlations. We then calculated each individual’s fitness based on a Gaussian surface (4):

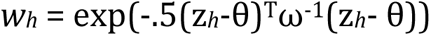

where *w_h_* is the fitness of individual *h*, *z_h_* is a vector of the observed phenotypic values for individual *h*, *ω* is the selection surface, and θ is the optima for traits (0). Truncation selection was applied based on fitness, with the 50% of individuals possessing the highest fitness surviving (main results). In an additional set of simulations, stronger truncation selection was applied and only 10% of the population survived.

Following selection in either framework, the next generation was constructed using an individual variance components approach (1). Specifically, the next generation was generated as described above based on the trait means, variances and covariances of survivors. Selection therefore acted via changes in means and variances and drift during the selection simulations was due to sampling error from the selection shaped phenotypic distributions.

##### Drift alone

For populations evolving via drift alone phenotypes were generated as for Gaussian adaptive landscapes. Composition of subsequent generations was likewise generated based on the means and variances of the prior generation, without selection. The drift model therefore was simply a model of sampling error.

Each of five modeling scenarios (simple landscapes, drift alone, three Holey landscapes with *p* = 0.2, 0.5, or 0.8) was simulated 250 times for populations of 7500 individuals and for 100 generations for each population. All modeling code is available at https://github.com/DochtermannLab/Wright_vs_Holey.

##### Statistical Comparison of Evolutionary Metrics

To clarify differences in evolutionary outcomes across modeling scenarios, we summarized evolutionary outcomes at the level of **G** matrices based on several metrics:

1. *λ*_2_⁄*λ*_1_; results for this metric are presented in the main text
2. *λ*_1_⁄∑ *λ*; this is a commonly used summary value and represents the proportion of variation captured by dominant eigenvalue. This can be interpreted as the proportion variation in the main dimension of covariance
3. ∑ *λ*; matrix trace, the total variation present. For simulations this is informative as to whether a particular process results in the loss of more or less variation
4. ē: average evolvability across dimensions (5). Evolutionary potential throughout multivariate space
5. ā: average reduction in evolvability due to trait covariance (5). Can be interpreted as how constrained evolutionary responses are based on correlations. At the extreme, an average autonomy of 0 would indicate absolute constraints on responses to selection and an average autonomy of 1 indicates evolutionary independence. Values between 0 and 1 represent quantitative constraints.

We compared these metrics across drift, Gaussian, and holey landscape simulations, following the main text, based on ANOVA followed by post-hoc comparisons based on calculation of Tukey’s Honest Significant Differences (HSD).

##### Post-hoc Parameter Exploration

The above modeling scenarios were used for our overall general analyses and for comparison to observed values. However, to explore whether our modeling outcomes were due to fundamentally different and generalizable outcomes or instead emerged from peculiarities of initial parameters, we expanded our analyses in two ways.

First, in addition to the moderate/weak strength of truncation selection modeled above (0.5), we also modeled stronger selection where only 10% of individuals survived. For this stronger strength of selection we again conducted 250 simulations of 7500 individuals for 100 generations. These simulations were included in the above analyses.

Second, to more broadly examine the sensitivity of our results to different starting values, we conducted simulation studies for our selection model, our model of drift, and our model of evolution on flat holey landscapes. For each modeling scenario (Gaussian surfaces, drift, Holey landscapes) we conducted 1000 simulations where both the magnitude of initial genetic variation in each trait varied and h^2^ varied (h^2^ was defined independently). For each scenario we then explored how other changes in starting parameters affected the eigenstructure of **G** (Table S2).

We then quantitatively assessed the relevance of each varied parameter on *λ*_2_⁄*λ*_1_— within modeling scenario—using linear models. All two-way interactions were included in analyses and variables (model parameters) were mean centered but unscaled. We then qualitatively compared *λ*_2_⁄*λ*_1_ across modeling scenarios based on heat plots.

**Table S2.**
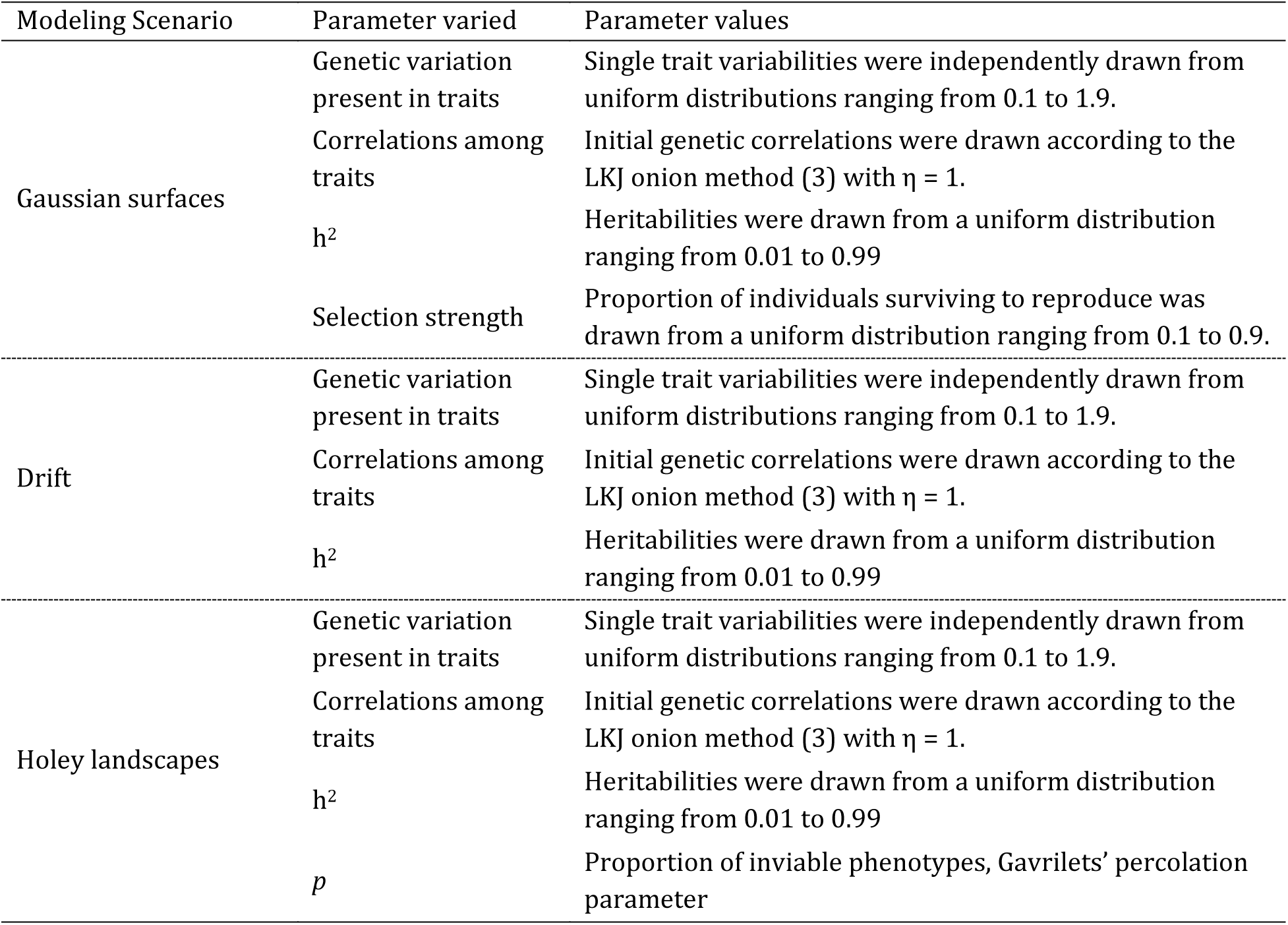
Parameters varied across simulation iterations by modeling scenario and range of possible values

#### Empirically Estimated **G** Matrices

##### Observed patterns of multivariate genetic variation

We conducted a literature review with Web of Science to search the journals American Naturalist, Ecology and Evolution, Evolution, Evolutionary Applications, Evolutionary Ecology, Genetics, Heredity, Journal of Evolutionary Biology, Journal of Heredity, Nature Ecology and Evolution, and the Proceedings of the Royal Society (B). These journals were searched using the terms “G matrix” on 14 May 2019, yielding a total of 272 articles. Each article was reviewed to determine if the article met inclusion criteria. Our inclusion criteria were:

1. A **G** matrix must have been estimated for more than 2 traits (i.e. > 2 × 2)
2. Must have been reported as variances and covariances (i.e. not genetic correlations)
3. Must not have been estimated for humans.

Based on these inclusion criteria, we ended up with 181 estimated **G** matrices (Fig S3). For each published **G** matrix, we calculated *λ*_2_⁄*λ*_1_ using a purpose-built R Shiny App ([link](https://dochtermannlab.shinyapps.io/matrixmetricsapp/)).

For each estimate we recorded the paper from which it was drawn (recorded as a unique study ID), taxonomic information (Kingdom through species epithet), trait category (life-history, physiology, morphology, behavior or mixed), the number of traits in the matrix, *λ*_1_, *λ*_2_, *λ*_2_⁄*λ*_1_, number of dimensions (6), number of dimensions divided by the number of traits, and all bibliographic information.

##### Phylogenetic Signal in *λ*_2_⁄*λ*_1_

To test for phylogenetic signal we fit a simple taxonomic mixed-effects model. This modeling approach incorporates the hierarchical non-independence due to taxonomic relationships but does not require a full phylogeny (7). Essentially, at each node of a phylogeny, relationships are modeled according to a star relationship. Each taxonomic grouping was included as a random effect, as was study ID, and the resulting model fit with the `lme4` package in R (8). From this model we estimated phylogenetic signal as the proportion of variation attributable to taxonomy, the variation attributable to study ID, and the residual variance. Confidence intervals were then estimated based on likelihood profile likelihoods.

##### Comparison of Observed Results to Simulation Results

Finally, we compared the observed values to the average for each of the simulation using the intercept coefficient of the above linear model. For this, t was calculated as (9):

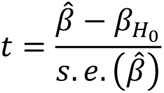

where 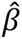 was the estimated intercept from the taxonomic model (above) and *β*_*H*0_ was a simulation average. p was calculated with degrees of freedom estimated using Satterthwaite’s method (df = 17.275).

### Supplemental Results

#### Simulation Models

##### Statistical Comparison of Evolutionary Metrics

Populations that evolved on different landscapes (drift alone, Gaussian, or holey) significantly differed from each other in the structure of **G** after 100 generations (Tables S3 – S7). Holey landscapes were characterized by a compression of most variation into the dominant dimension in multivariate space (Tables S3 & S4; Figures 2 & S5). Populations evolving on Gaussian landscapes were characterized by a drastic reduction in the total variation present, which was also reflected in reduced evolvability (Tables S6 & S7; Figures S5 & S6). The combination of high standing genetic variation and this variation being distributed across dimensions led to populations that evolved solely due to drift to exhibit significantly greater autonomy than observed in any of the other modeling scenarios (Table S7; Figure S8). This greater constraint in populations evolving on either Gaussian or holey landscapes is likely due to the loss of variation for populations evolving on Gaussian landscapes (Figures S6 & S7) and the compression of variation for populations evolving on holey landscapes (Figures 2 & S5).

**Table S3.** ANOVA and Tukey HSD results for *λ*_2_⁄*λ*_1_. Significantly greater genetic variation was maintained across all dimensions when populations evolved on Gaussian landscapes or due to drift than when evolving on holey landscapes (Figure 2, main text).

| ANOVA Results |  |  |  |  |  |
| --- | --- | --- | --- | --- | --- |
|  | df | SS | MSS | F | p |
| Simulation type | 5 | 54.98 | 10.996 | 343.5 | <0.01 |
| Residual | 1494 | 47.82 | 0.032 |  |  |
| Tukey HSD |  |  |  |  |  |
| Simulation Comparison | Difference | Lower | Upper | p |  |
| Holey p = 0.5-Holey p = 0.2 | -0.026 | -0.071 | 0.020 | 0.589 |  |
| Holey p = 0.8-Holey p = 0.2 | 0.011 | -0.035 | 0.057 | 0.984 |  |
| Wright 0.1-Holey p = 0.2 | 0.293 | 0.248 | 0.339 | <0.01 |  |
| Wright 0.5-Holey p = 0.2 | 0.374 | 0.329 | 0.420 | <0.01 |  |
| Drift-Holey p = 0.2 | 0.437 | 0.391 | 0.483 | <0.01 |  |
| Holey p = 0.8-Holey p = 0.5 | 0.037 | -0.009 | 0.082 | 0.198 |  |
| Wright 0.1-Holey p = 0.5 | 0.319 | 0.273 | 0.365 | <0.01 |  |
| Wright 0.5-Holey p = 0.5 | 0.400 | 0.354 | 0.446 | <0.01 |  |
| Drift-Holey p = 0.5 | 0.463 | 0.417 | 0.508 | <0.01 |  |
| Wright 0.1-Holey p = 0.8 | 0.282 | 0.237 | 0.328 | <0.01 |  |
| Wright 0.5-Holey p = 0.8 | 0.363 | 0.318 | 0.409 | <0.01 |  |
| Drift-Holey p = 0.8 | 0.426 | 0.380 | 0.472 | <0.01 |  |
| Wright 0.5-Wright 0.1 | 0.081 | 0.035 | 0.127 | <0.01 |  |
| Drift-Wright 0.1 | 0.144 | 0.098 | 0.189 | <0.01 |  |
| Drift-Wright 0.5 | 0.063 | 0.017 | 0.108 | <0.01 |  |

**Table S4.**
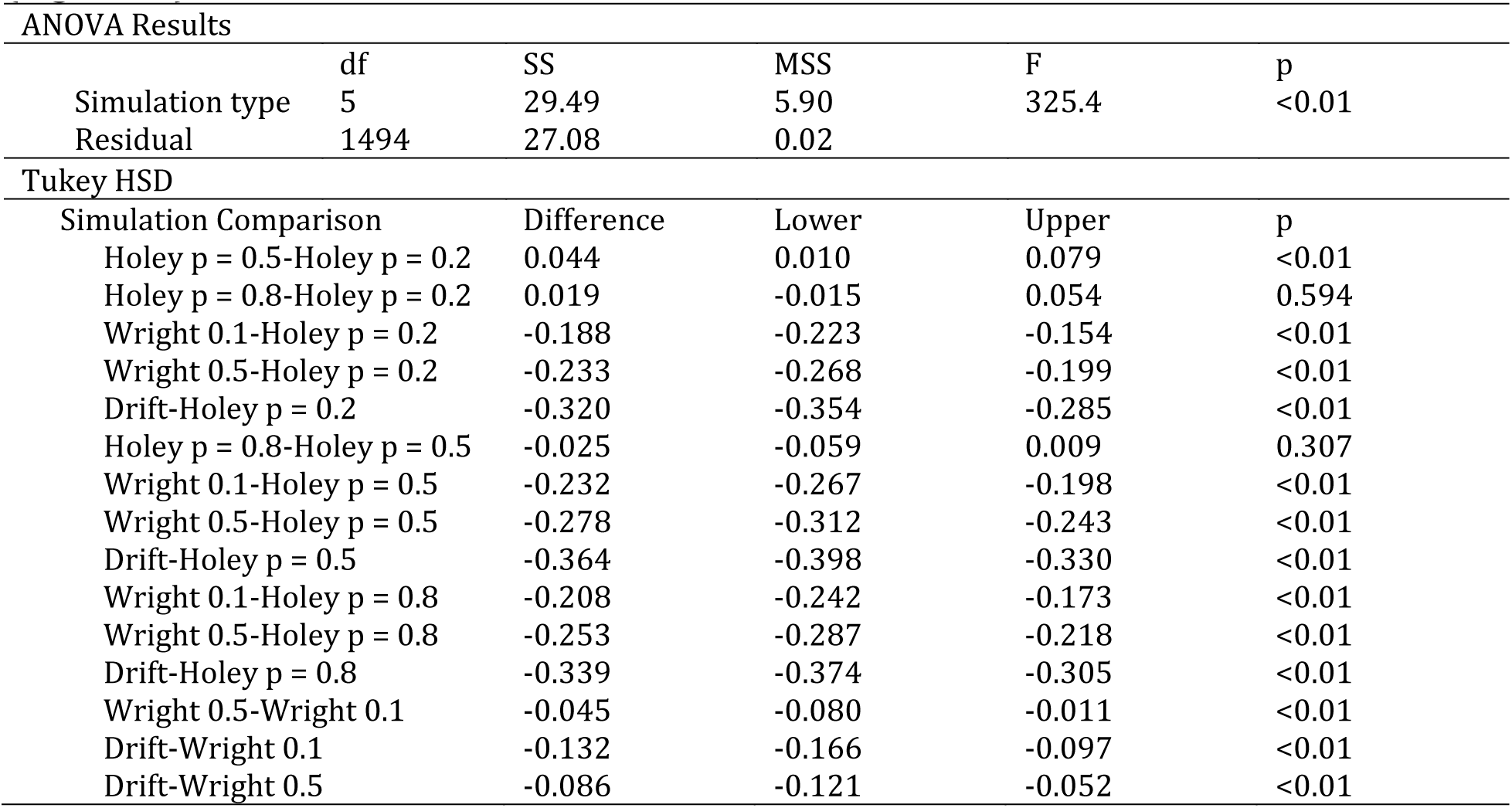
ANOVA and Tukey HSD results for *λ*_1_⁄∑ *λ*. Significantly greater proportional genetic variation was retained in the dominant multivariate direction for populations that evolved on Gaussian landscapes or via drift than when evolving on holey landscapes (Figure S5).

**Table S5.**
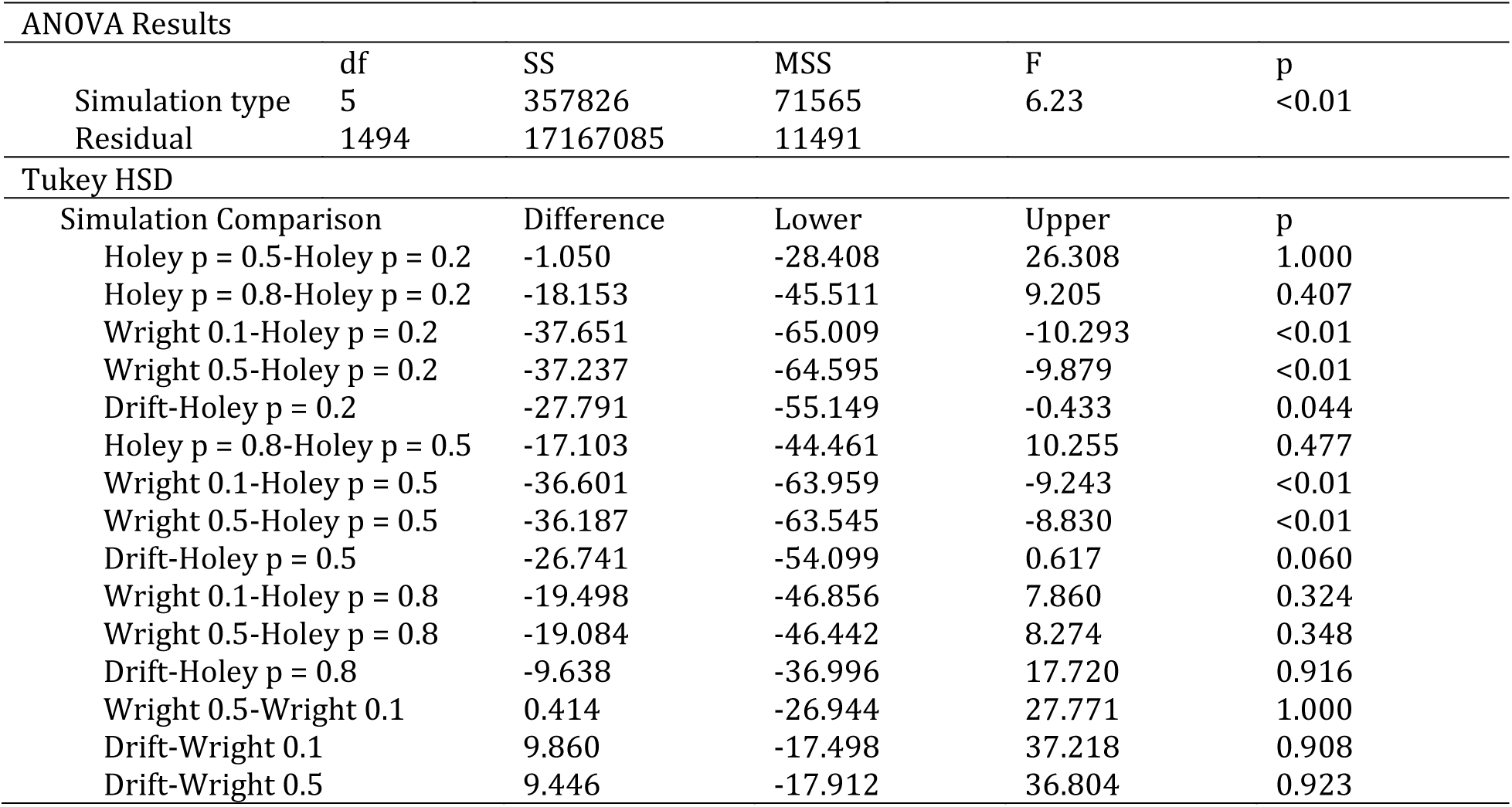
ANOVA and Tukey HSD results for the total genetic variation in populations at the end of simulations ∑ *λ*. The amount of total variation significantly varied across simulation types. Populations that evolved on Gaussian landscapes lost considerably more genetic variation than those evolving on other landscapes (Figure S6).

**Table S6.** ANOVA and Tukey HSD results for evolvability, ē. Because more genetic variation was maintained when populations evolved on holey landscapes or drift (Table S5), evolvability was significantly lower when populations evolved on Gaussian landscapes (Figure S7). (evolvability is just the matrix trace divided by the number of traits)

| ANOVA Results |  |  |  |  |  |
| --- | --- | --- | --- | --- | --- |
|  | df | SS | MSS | F | p |
| Simulation type | 5 | 3578 | 715.7 | 6.23 | <0.01 |
| Residual | 1494 | 171671 | 114.9 |  |  |
| Tukey HSD |  |  |  |  |  |
| Simulation Comparison | Difference | Lower | Upper | p |  |
| Holey p = 0.5-Holey p = 0.2 | -0.105 | -2.841 | 2.631 | 1.000 |  |
| Holey p = 0.8-Holey p = 0.2 | -1.815 | -4.551 | 0.921 | 0.407 |  |
| Wright 0.1-Holey p = 0.2 | -3.765 | -6.501 | -1.029 | <0.01 |  |
| Wright 0.5-Holey p = 0.2 | -3.724 | -6.460 | -0.988 | <0.01 |  |
| Drift-Holey p = 0.2 | -2.779 | -5.515 | -0.043 | 0.044 |  |
| Holey p = 0.8-Holey p = 0.5 | -1.710 | -4.446 | 1.025 | 0.477 |  |
| Wright 0.1-Holey p = 0.5 | -3.660 | -6.396 | -0.924 | <0.01 |  |
| Wright 0.5-Holey p = 0.5 | -3.619 | -6.355 | -0.883 | <0.01 |  |
| Drift-Holey p = 0.5 | -2.674 | -5.410 | 0.062 | 0.060 |  |
| Wright 0.1-Holey p = 0.8 | -1.950 | -4.686 | 0.786 | 0.324 |  |
| Wright 0.5-Holey p = 0.8 | -1.908 | -4.644 | 0.827 | 0.348 |  |
| Drift-Holey p = 0.8 | -0.964 | -3.700 | 1.772 | 0.916 |  |
| Wright 0.5-Wright 0.1 | 0.041 | -2.694 | 2.777 | 1.000 |  |
| Drift-Wright 0.1 | 0.986 | -1.750 | 3.722 | 0.908 |  |
| Drift-Wright 0.5 | 0.945 | -1.791 | 3.680 | 0.923 |  |

**Table S7.** ANOVA and Tukey HSD results for autonomy, ā. Significantly greater variation was maintained across all dimensions when populations evolved on Gaussian landscapes or due to drift than when evolving on holey landscapes (Figure S8).

| ANOVA Results |  |  |  |  |  |
| --- | --- | --- | --- | --- | --- |
|  | df | SS | MSS | F | p |
| Simulation type | 5 | 43.61 | 8.72 | 518.3 | <0.01 |
| Residual | 1494 | 25.14 | 0.02 |  |  |
| Tukey HSD |  |  |  |  |  |
| Simulation Comparison | Difference | Lower | Upper | p |  |
| Holey p = 0.5-Holey p = 0.2 | -0.021 | -0.054 | 0.012 | 0.479 |  |
| Holey p = 0.8-Holey p = 0.2 | -0.042 | -0.075 | -0.009 | <0.01 |  |
| Wright 0.1-Holey p = 0.2 | -0.148 | -0.181 | -0.115 | <0.01 |  |
| Wright 0.5-Holey p = 0.2 | 0.395 | 0.362 | 0.428 | <0.01 |  |
| Drift-Holey p = 0.2 | 0.077 | 0.044 | 0.110 | <0.01 |  |
| Holey p = 0.8-Holey p = 0.5 | -0.022 | -0.055 | 0.011 | 0.424 |  |
| Wright 0.1-Holey p = 0.5 | -0.127 | -0.160 | -0.094 | <0.01 |  |
| Wright 0.5-Holey p = 0.5 | 0.415 | 0.382 | 0.448 | <0.01 |  |
| Drift-Holey p = 0.5 | 0.098 | 0.064 | 0.131 | <0.01 |  |
| Wright 0.1-Holey p = 0.8 | -0.106 | -0.139 | -0.072 | <0.01 |  |
| Wright 0.5-Holey p = 0.8 | 0.437 | 0.404 | 0.470 | <0.01 |  |
| Drift-Holey p = 0.8 | 0.119 | 0.086 | 0.152 | <0.01 |  |
| Wright 0.5-Wright 0.1 | 0.543 | 0.509 | 0.576 | <0.01 |  |
| Drift-Wright 0.1 | 0.225 | 0.192 | 0.258 | <0.01 |  |
| Drift-Wright 0.5 | -0.318 | -0.351 | -0.285 | <0.01 |  |

##### Post-hoc Parameter Exploration

For populations evolving on Gaussian landscapes, compression of genetic variation into the leading dimension decreased with increasing heritability and an increasing strength of selection (Table S8, Figure S9). No two-way interaction was statistically significant. Put another way, *λ*_2_⁄*λ*_1_, increased with heritability and the strength of selection and average *λ*_2_⁄*λ*_1_ was 0.68 for average parameter values (Table S8).

For populations evolving solely due to drift, *λ*_2_⁄*λ*_1_ increased with greater initial total genetic variation (Table S9). However, the strength of this effect was minimal. More dramatically, *λ*_2_⁄*λ*_1_ significantly and strongly decreased with increasing average initial absolute genetic correlation (Table S9). At the extreme, *λ*_2_⁄*λ*_1_ approached 0 as the average initial absolute correlation approaches 1. No two-way interaction was statistically significant. Average *λ*_2_⁄*λ*_1_ was 0.69 for average parameter values (Table S9).

When evolving on holey landscapes, and consistent with prior simulation comparisons, *λ*_2_⁄*λ*_1_ was lower for average parameter values (0.42, Table S10). Compression into a single dimension also increased with increasing heritability and increasing average absolute initial correlations (Table S10).

Genetic variation was more strongly compressed into a primary dimension when populations evolved on holey landscapes versus when they evolved due to drift or due to selection on Gaussian surfaces (Tables S8 – S10; Figures S9 – S11). This was a surprisingly robust result regardless of the starting parameters of a simulation (Figures S9 – S11). This parameter robustness (10) supports the generality of our modeling. Unfortunately, we were not able to investigate other forms of robustness (10) due to computational limitations.

Our parameter exploration approach covered a wide range of possible values, making results from this exploration generalizable (Figures S12 – S14).

**Table S8.**
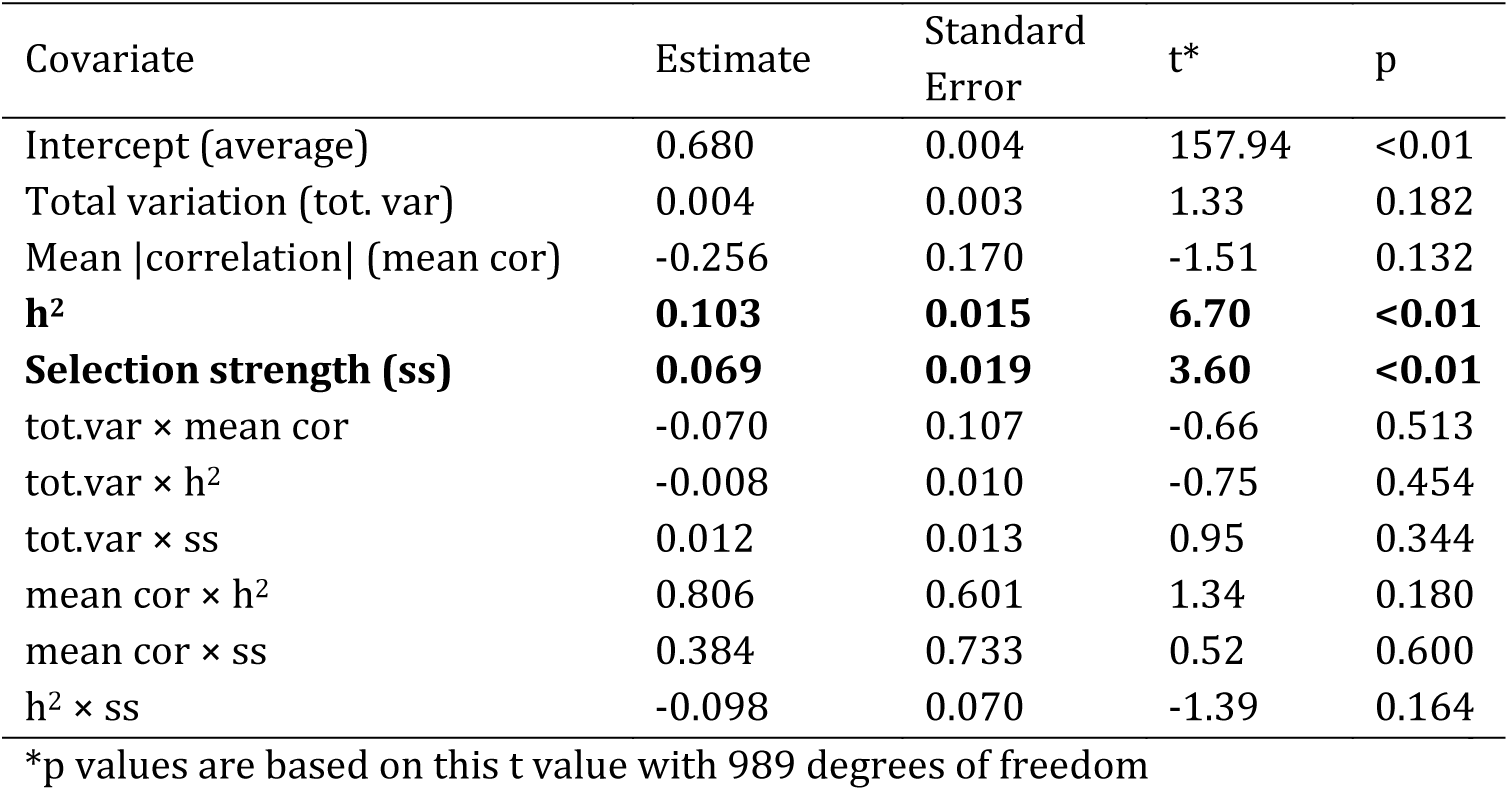
Linear modeling results for Gaussian landscape parameter exploration. All covariates were modeled while centered (but not variance standardized).

**Table S9.**
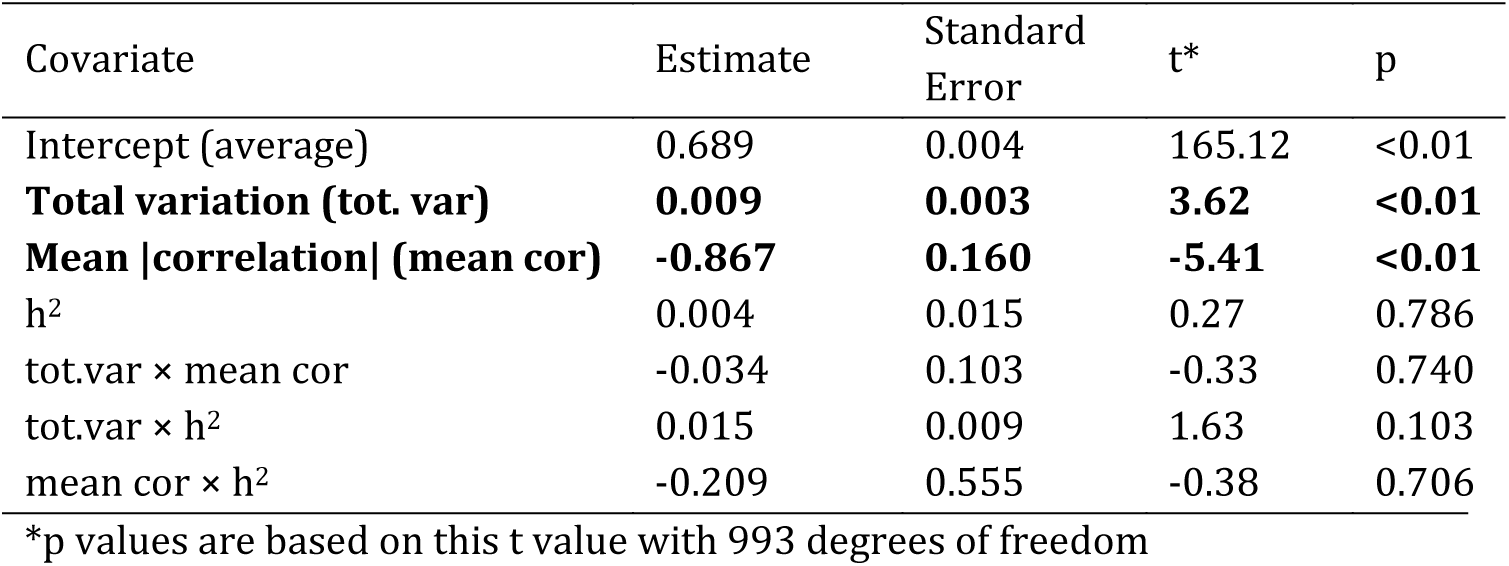
Linear modeling results for parameter exploration of the drift model. All covariates were modeled while centered (but not variance standardized).

**Table S8.**
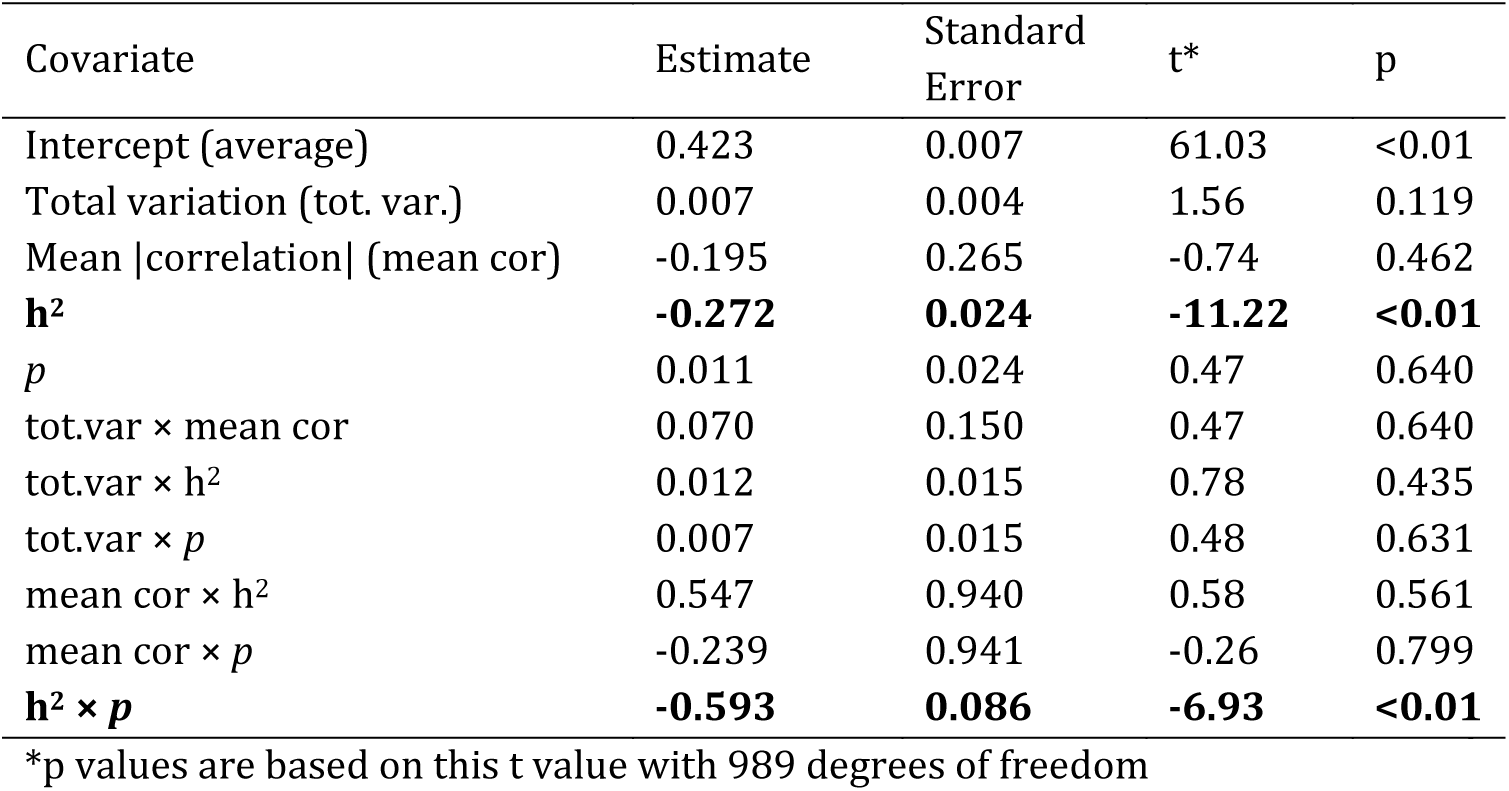
Linear modeling results for holey landscape parameter exploration. All covariates were modeled while centered (but not variance standardized).

#### Empirically Estimated G Matrices

##### Phylogenetic Signal in λ_2_⁄λ_1_

**Table S9.** Variances for *λ*_2_⁄*λ*_1_—with associated 95% confidence intervals—at each taxonomic level, for study ID, and residual. Proportion of variation for taxonomy, study ID, and residual are also provided

| Variance component | Estimate (95% CI) | Proportion of variance |
| --- | --- | --- |
| Study ID | 0.026 (0.013 : 0.048) | 0.45 |
| Taxonomy | 0.003 | 0.05 |
| species | 0 (0 : 0.01) |  |
| Genus | 0 (0 : 0.016) |  |
| Family | 0.003 (0 : 0.02) |  |
| Order | 0 (0 : 0.018) |  |
| Class | 0 (0 : 0.008) |  |
| Phylum | 0 (0 : 0.007) |  |
| Kingdom | 0 (0 : 0.011) |  |
| Residual | 0.029 (0.023 : 0.037) | 0.50 |

##### Comparison of Observed Results to Simulation Results

Observed results did not significantly differ from simulated populations that evolved on holey landscapes (Figure 2; Table S10). This conclusion based on *λ*_2_⁄*λ*_1_ is consistent when values of *λ*_1_⁄∑ *λ* are compared between simulated and empirical datasets (Figure S15)

**Table S10.**
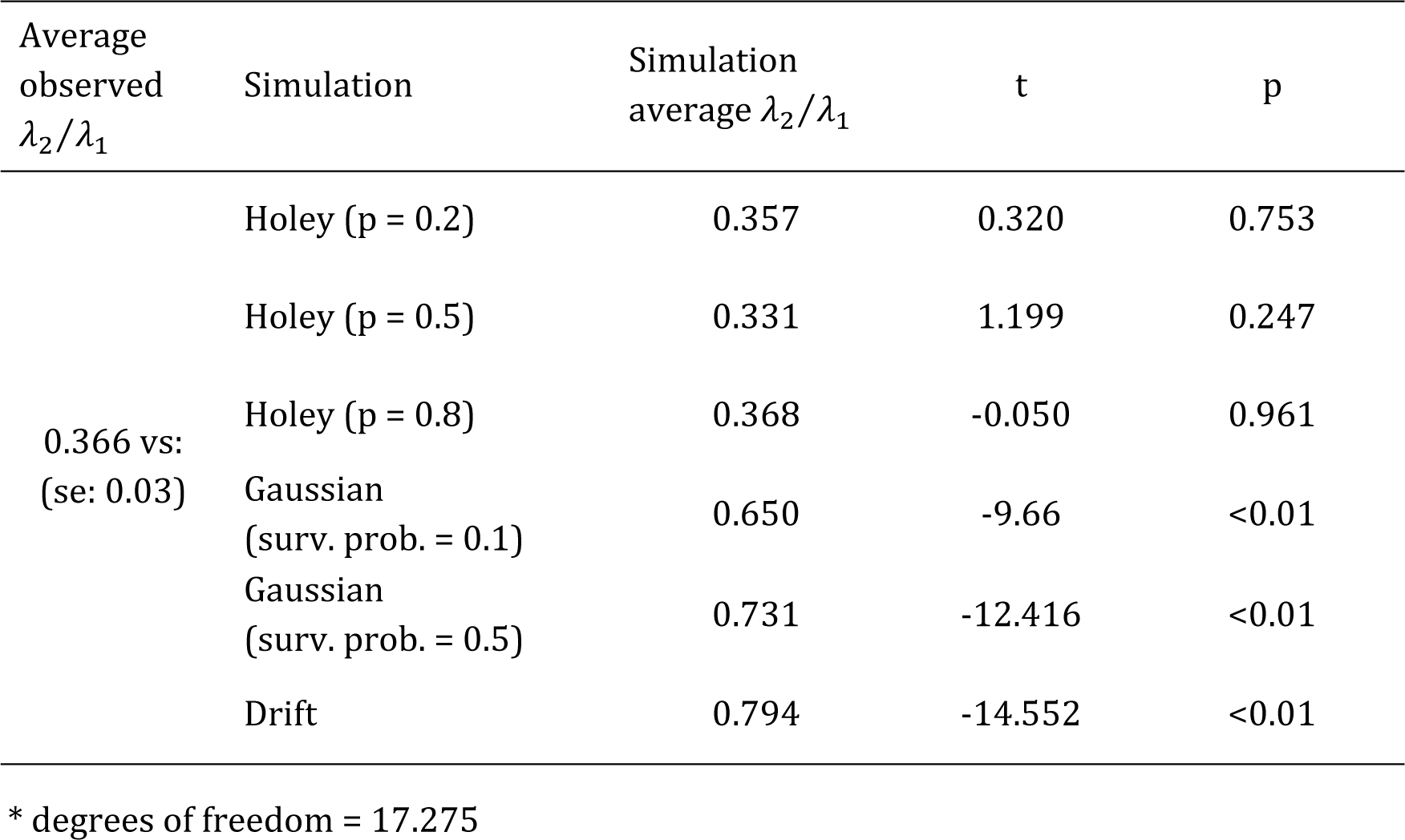
t values and associated p values for the comparison of the observed average of *λ*_2_⁄*λ*_1_ versus the average *λ*_2_⁄*λ*_1_ for each set of simulations. The observed average and its standard error was taken from a taxonomic mixed-effects model.

| Average<br>observed<br>$\lambda_2/\lambda_1$ | Simulation | Simulation<br>average $\lambda_2/\lambda_1$ | t | p |
| --- | --- | --- | --- | --- |
| 0.366 vs:<br>(se: 0.03) | Holey (p = 0.2) | 0.357 | 0.320 | 0.753 |
|  | Holey (p = 0.5) | 0.331 | 1.199 | 0.247 |
|  | Holey (p = 0.8) | 0.368 | -0.050 | 0.961 |
|  | Gaussian<br>(surv. prob. = 0.1) | 0.650 | -9.66 | <0.01 |
|  | Gaussian<br>(surv. prob. = 0.5) | 0.731 | -12.416 | <0.01 |
|  | Drift | 0.794 | -14.552 | <0.01 |

\* degrees of freedom = 17.275

**Figure S1.**
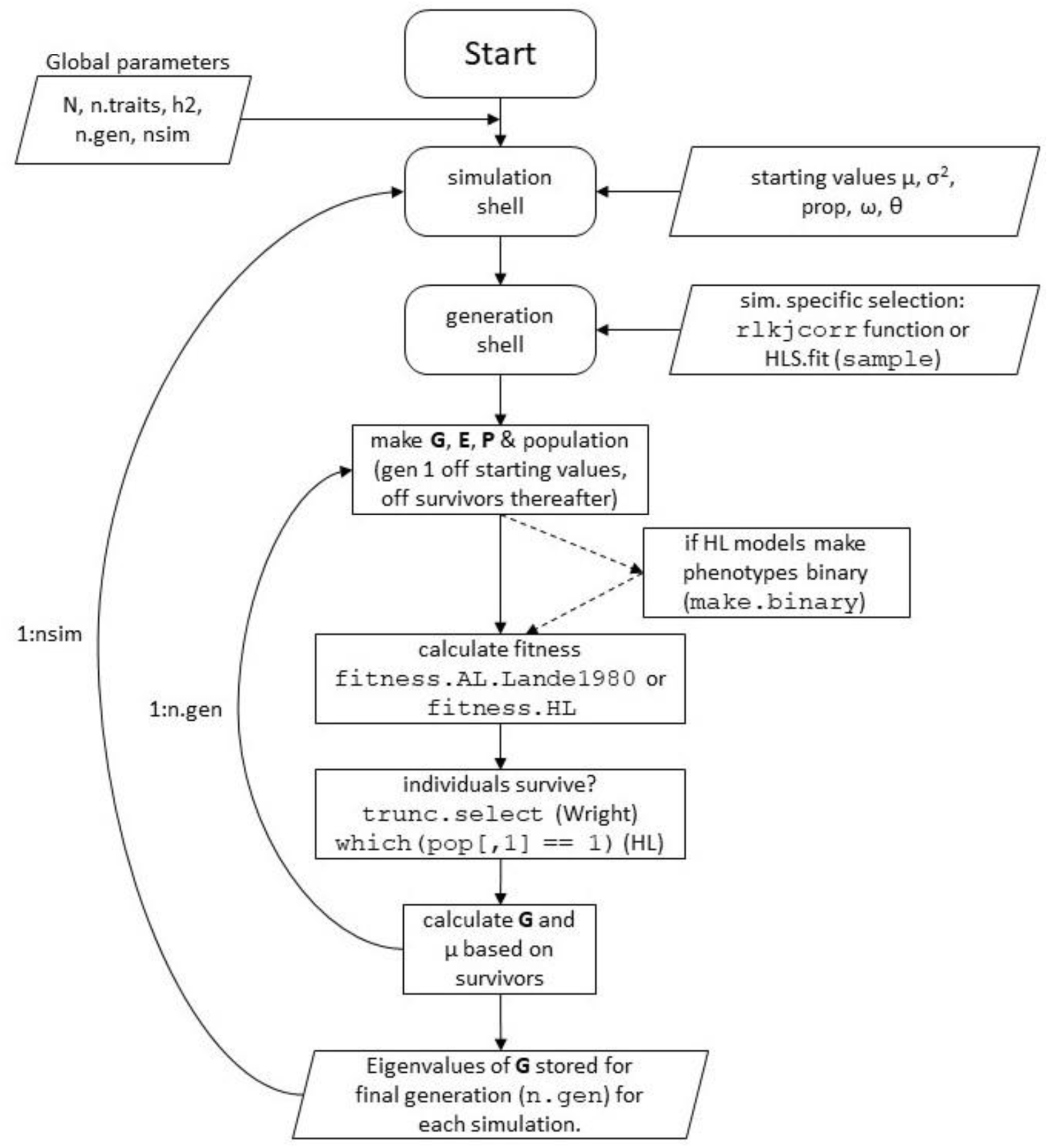
Model flow diagram for HL and gaussian landscapes

**Figure S2.**
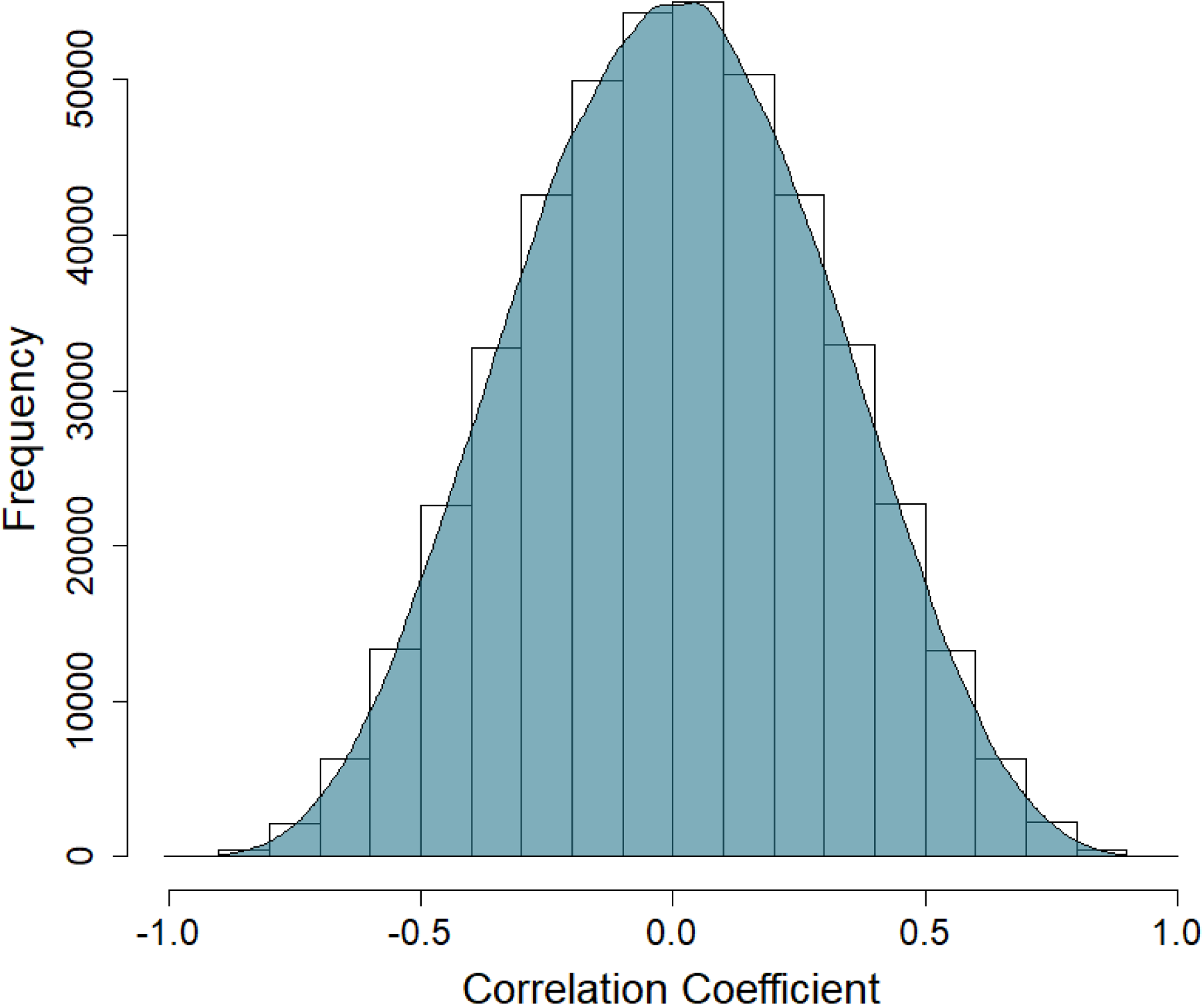
Distribution of 450000 random correlations generated using the LKJ Onion method with k = 10.

**Figure S3.**
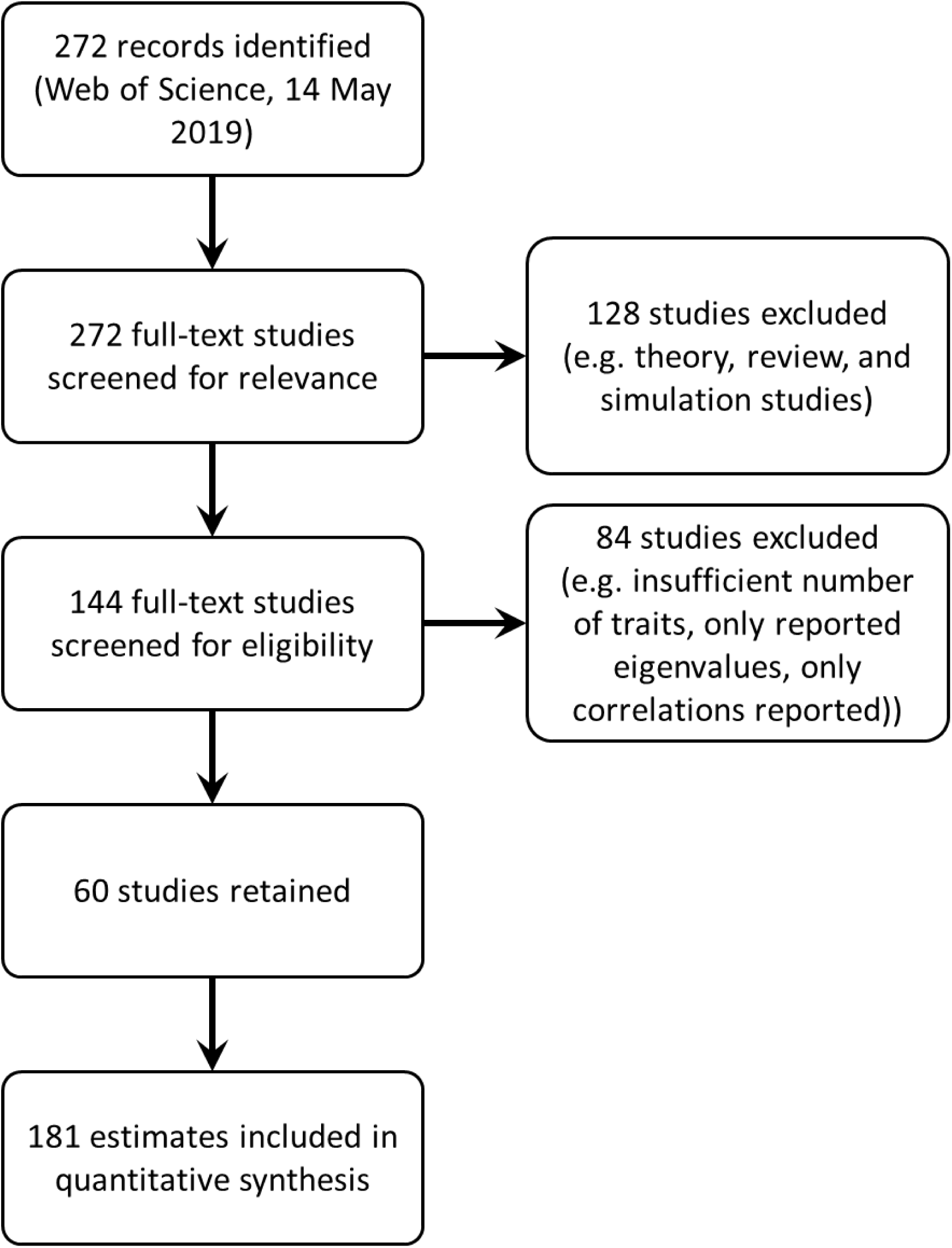
PRISMA diagram for studies and estimates included in taxonomic analyses.

**Figure S4.**
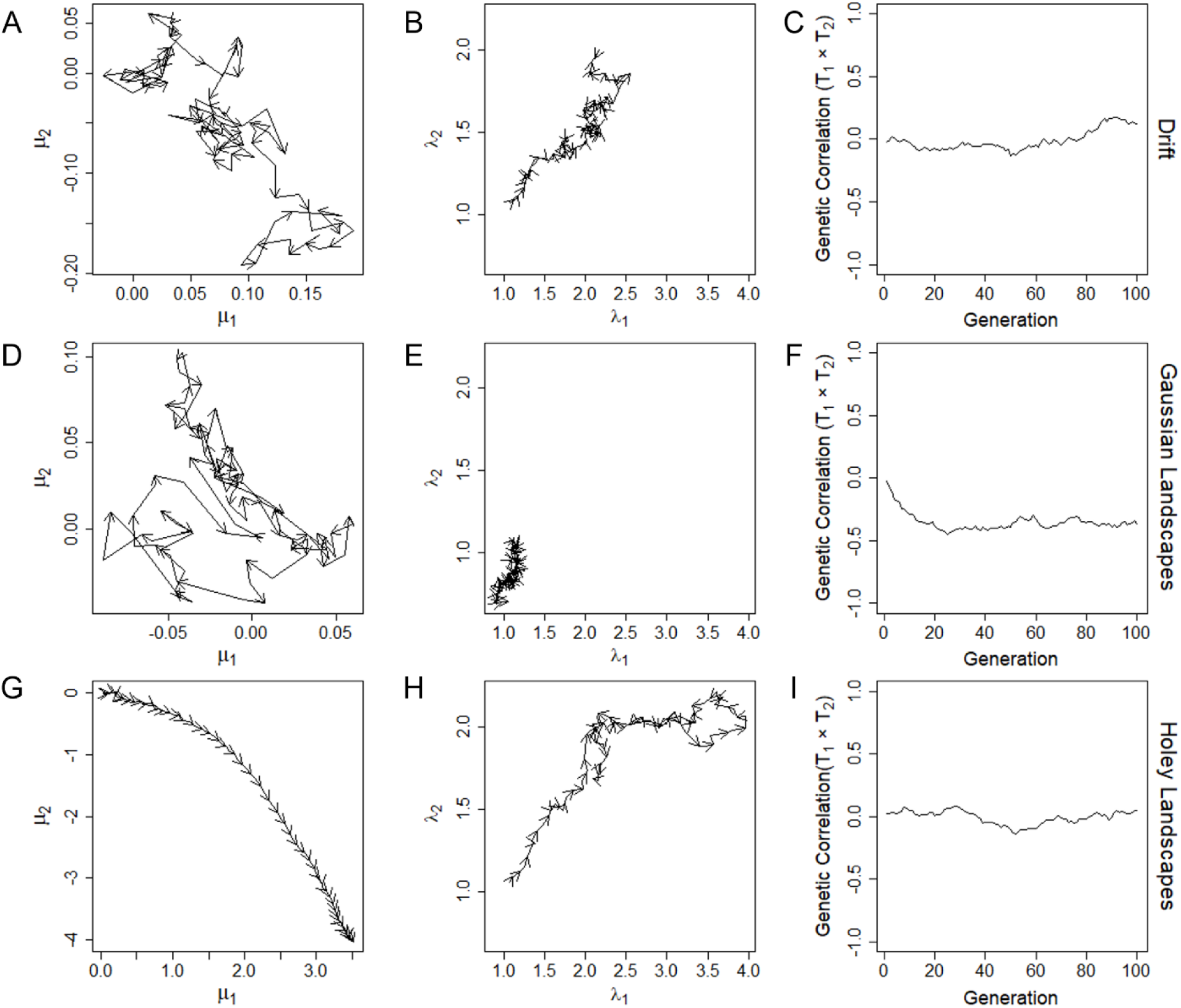
Single population comparisons of population evolution over 100 generations under drift (A – C), on a Gaussian landscape (D – F), and on a holey landscape (G – I). Arrow heads in A, B, D, E, G, and H indicate the direction of evolutionary change at every second generation. Evolutionary change in the average values for traits 1 and 2 (A, D, and G; note the different scales for axes) show little change for either drift and on a Gaussian landscape. In contrast, the population shows substantial and directional change in trait values on a holey landscape (G). This suggests the population is moving between holes in G but is restricted to a local optimum in D. The first and second eigenvalues (λ_1_ & λ_2_) show little total change due to drift (B), consistent decreases on a Gaussian landscape (E), and larger changes—including overall increases—on a holey landscape (H). This is consistent with the overall compression of variance reported elsewhere in the main and supplemental results. The bivariate genetic correlation between the first two traits shows little directional change under either drift of on holey landscapes (C and I) but rapid absolute increase followed by becoming static on a Gaussian landscape (F). As was the case for eigenvalues (E), this is consistent with stabilizing selection at a local optimum. These results are consistent across multiple runs, though exact trajectories vary and the sign of genetic correlations is equally likely to be positive as negative.

**Figure S5.**
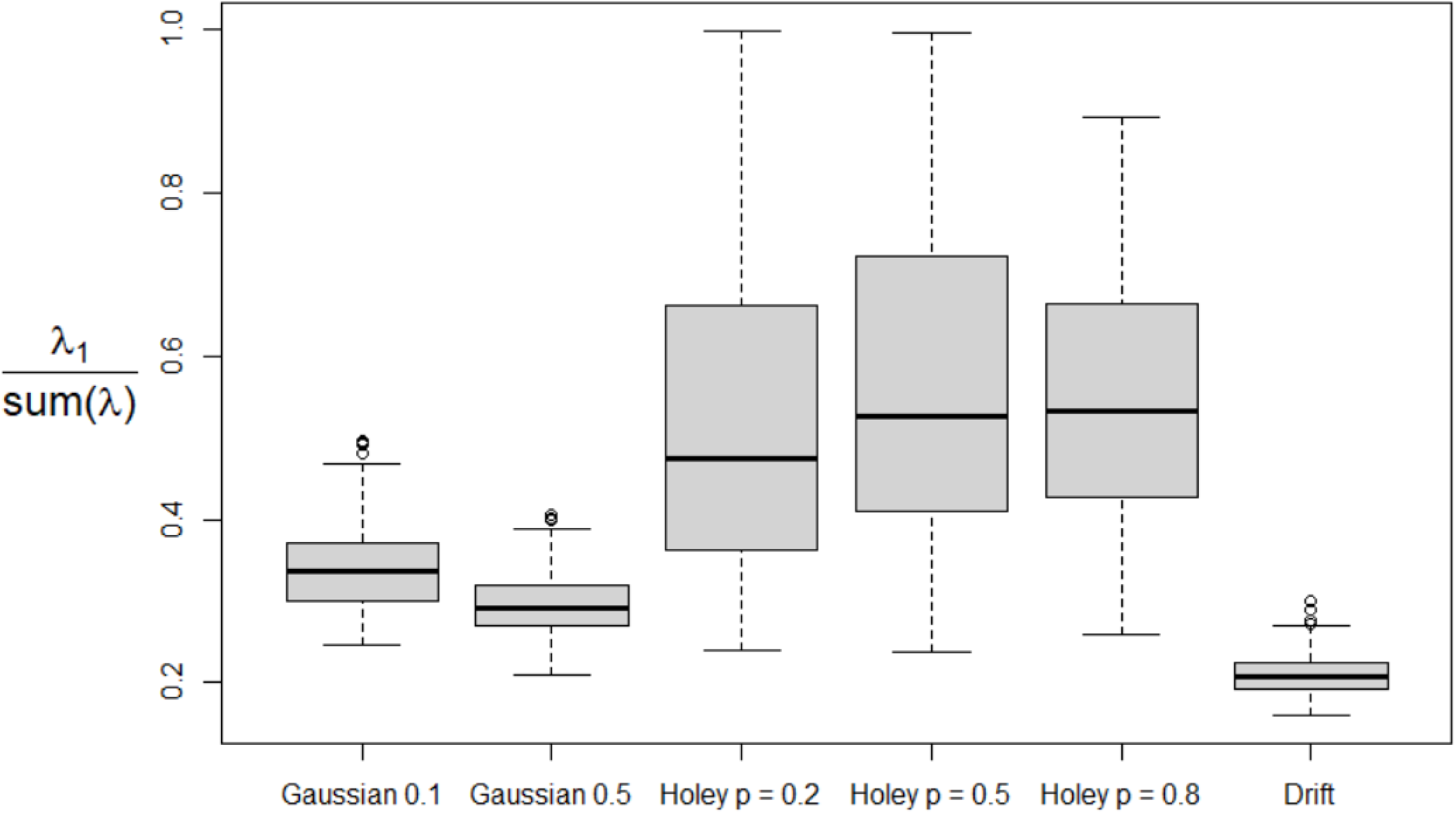
Variation was more evenly distributed across dimensions when populations evolved on Gaussian landscapes or due solely to drift. Consequently, less total variation was present in the first dimension (Table S4).

**Figure S6.**
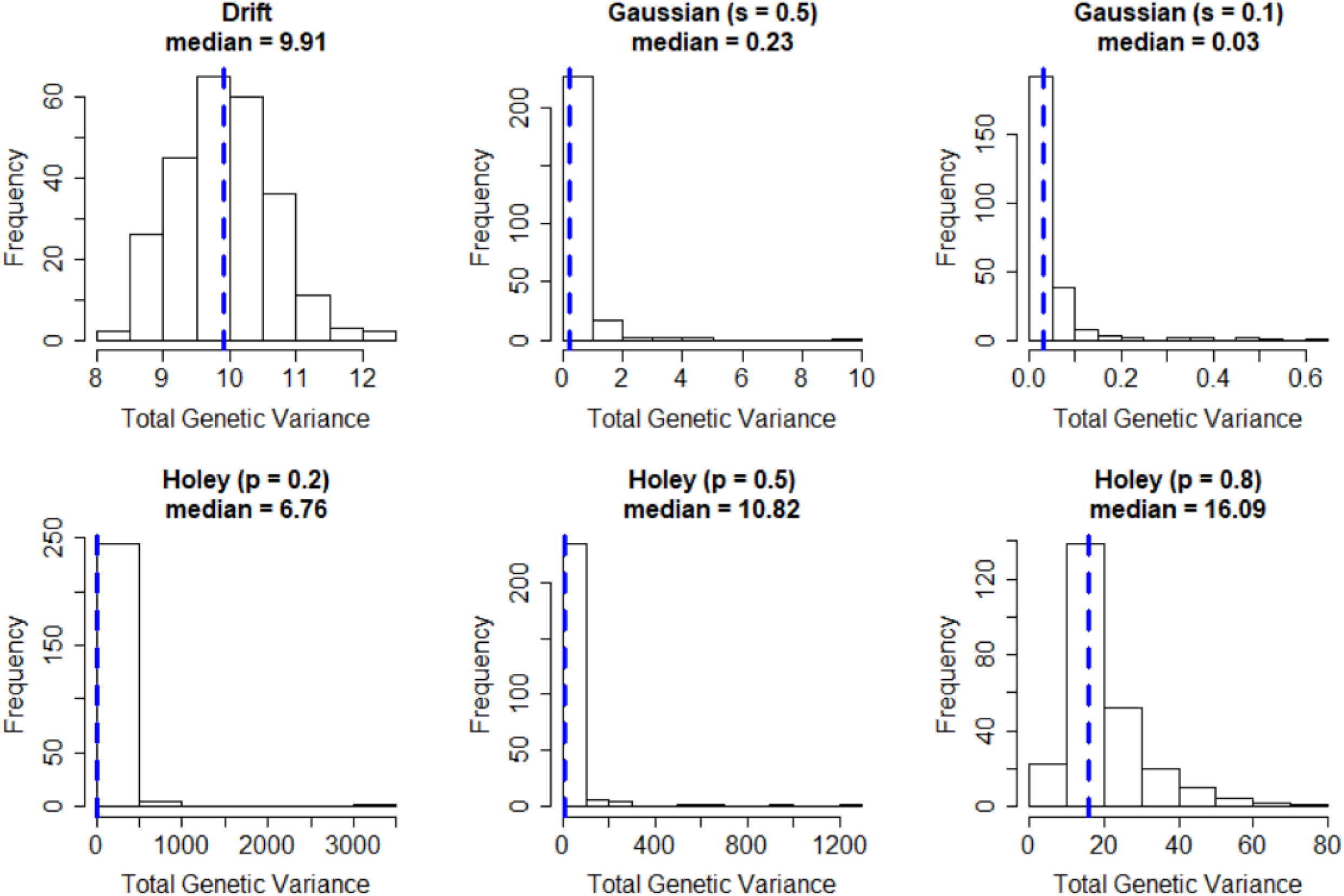
The total genetic variation present after 100 generations in each of six modeling conditions and across 250 simulations. Selection on Gaussian surfaces led to a significant reduction in the amount of variation present (Table S5). Dashed blue bars indicate the median value.

**Figure S7.**
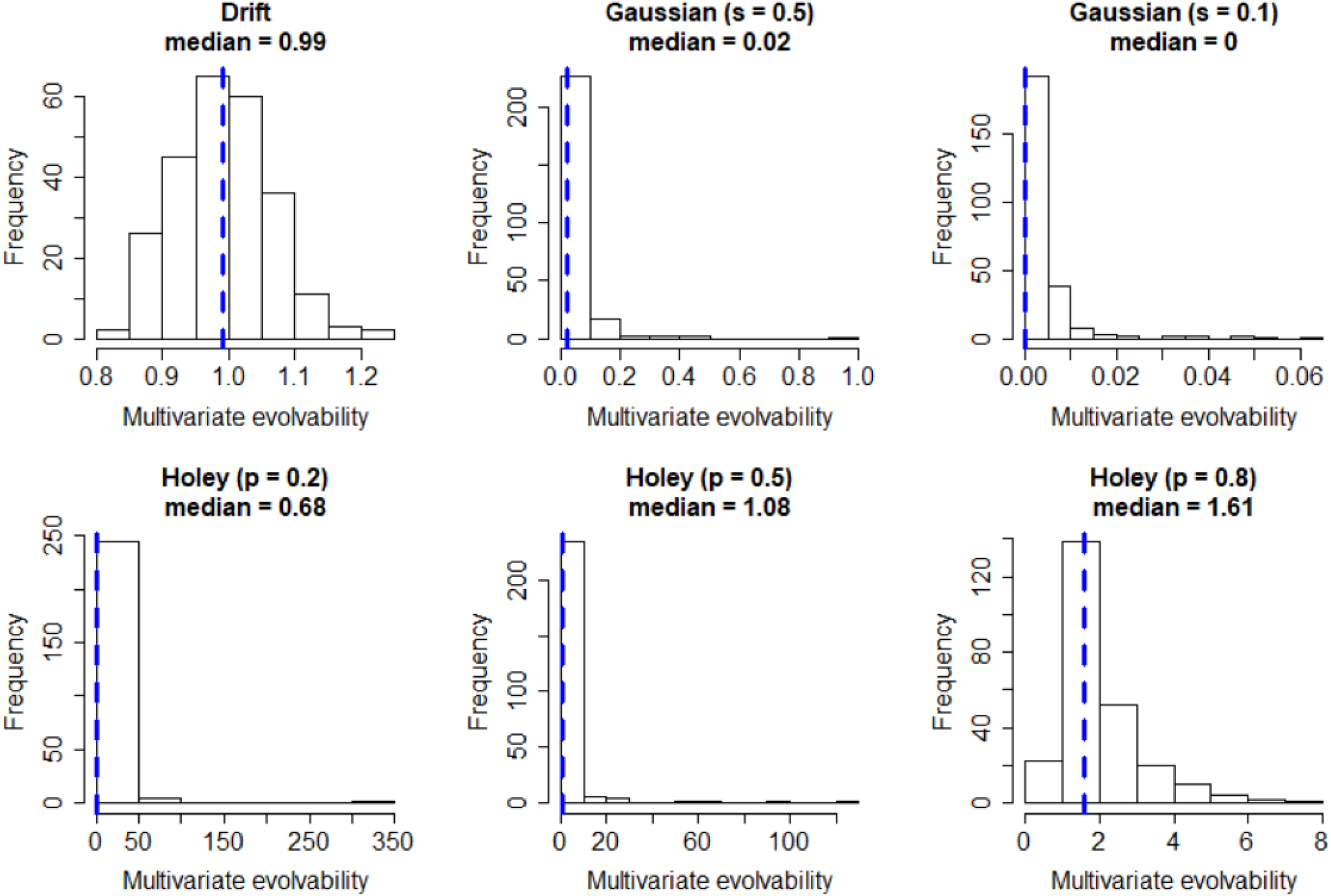
Multivariate evolvability after 100 generations in each of six modeling conditions and across 250 simulations. Selection on Gaussian surfaces led to a significant reduction in evolvability (Table S6). Dashed blue bars indicate the median value.

**Figure S8.**
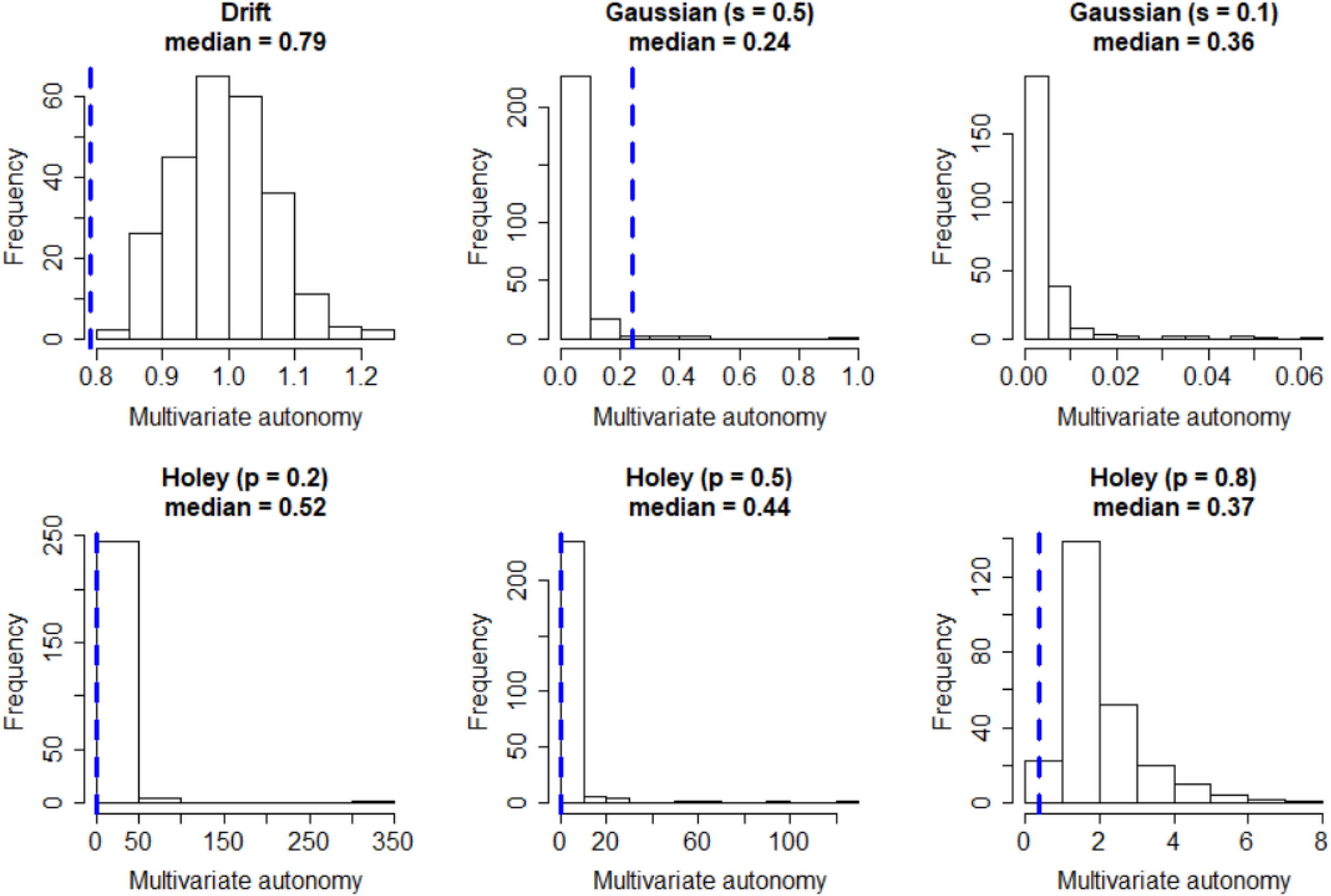
Multivariate autonomy after 100 generations in each of six modeling conditions and across 250 simulations. Selection on Gaussian surfaces led to a significant reduction in autonomy (Table S7). Dashed blue bars indicate the median value.

**Figure S9.**
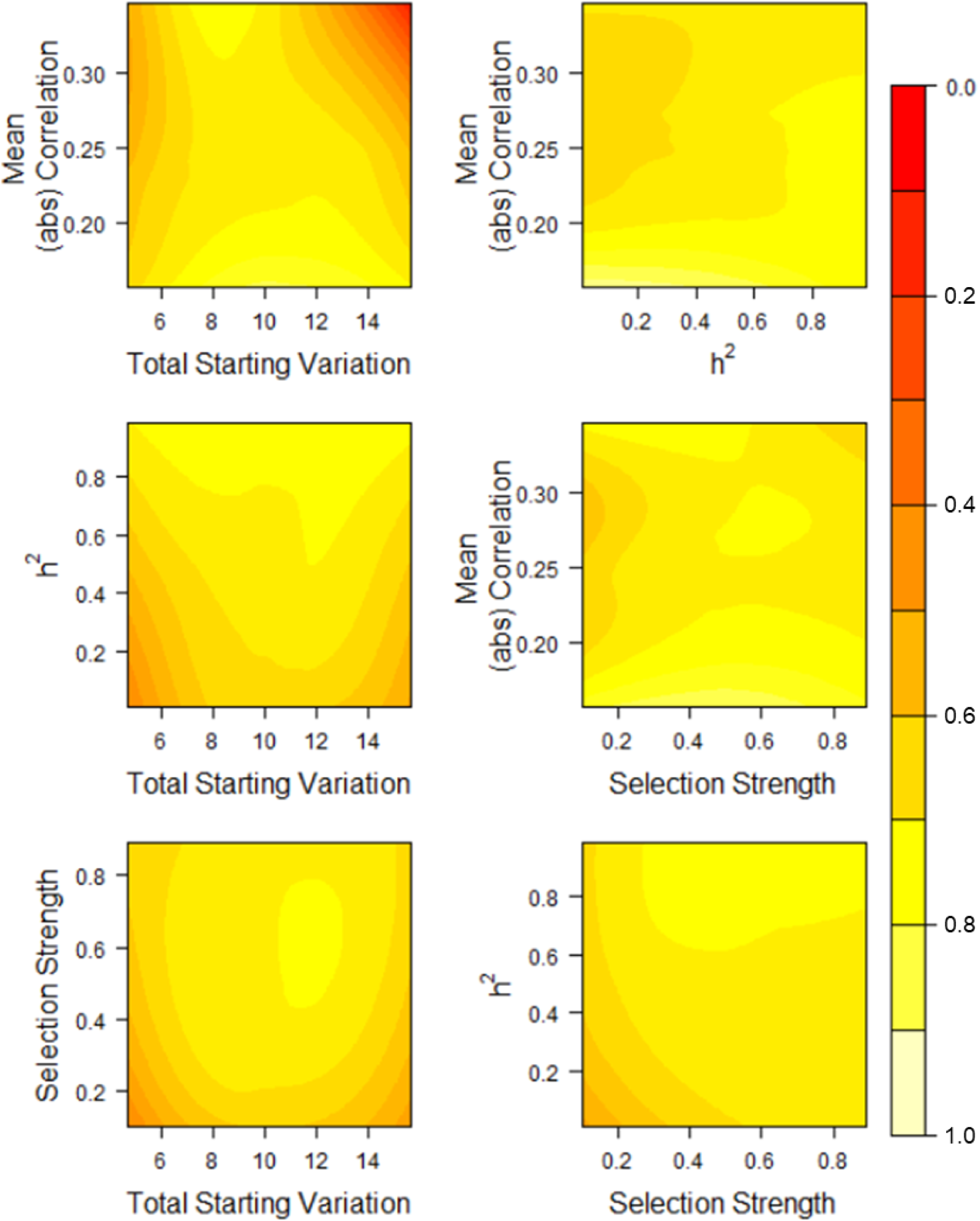
*λ*_2_⁄*λ*_1_ after selection on Gaussian surfaces remained high regardless of starting parameters (Table S8).

**Figure S10.**
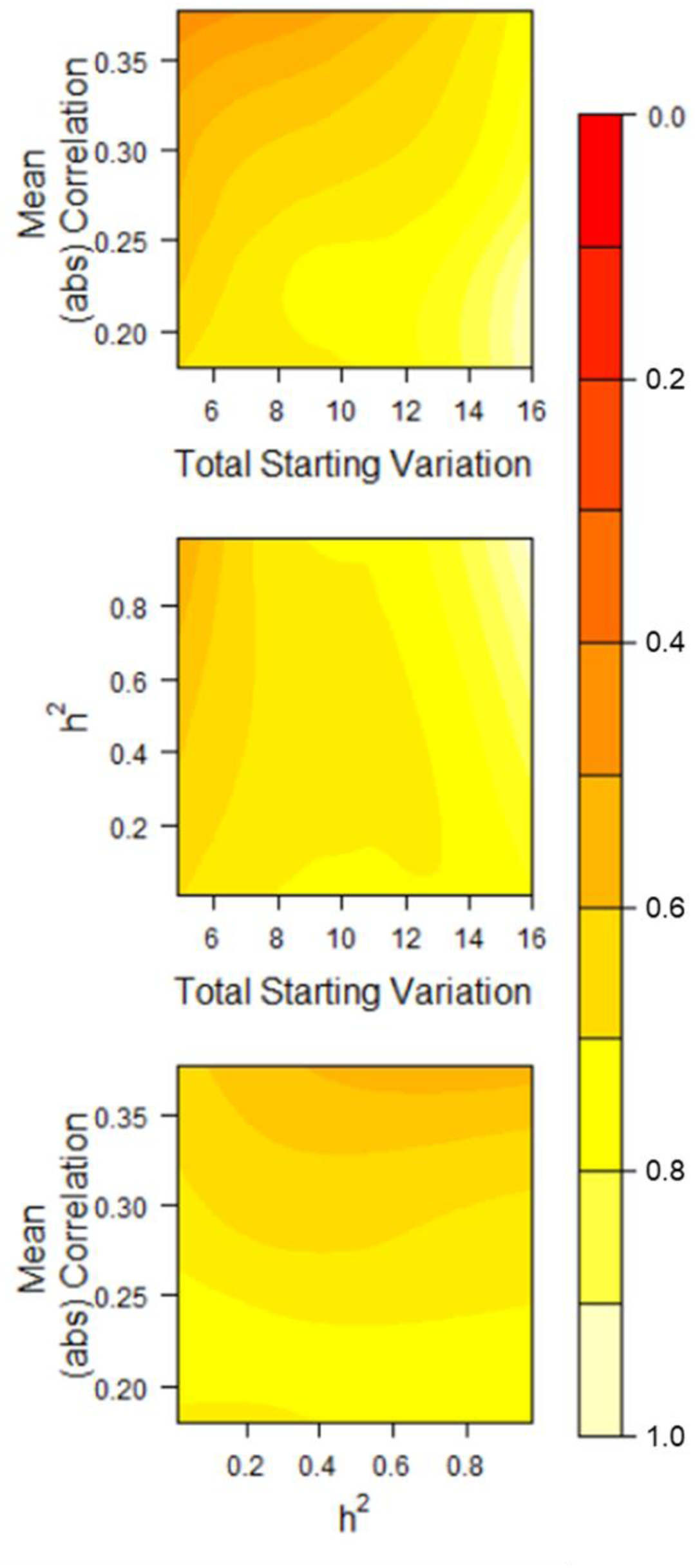
*λ*_2_⁄*λ*_1_ after evolution due to drift remained high regardless of starting parameters (Table S9).

**Figure S11.**
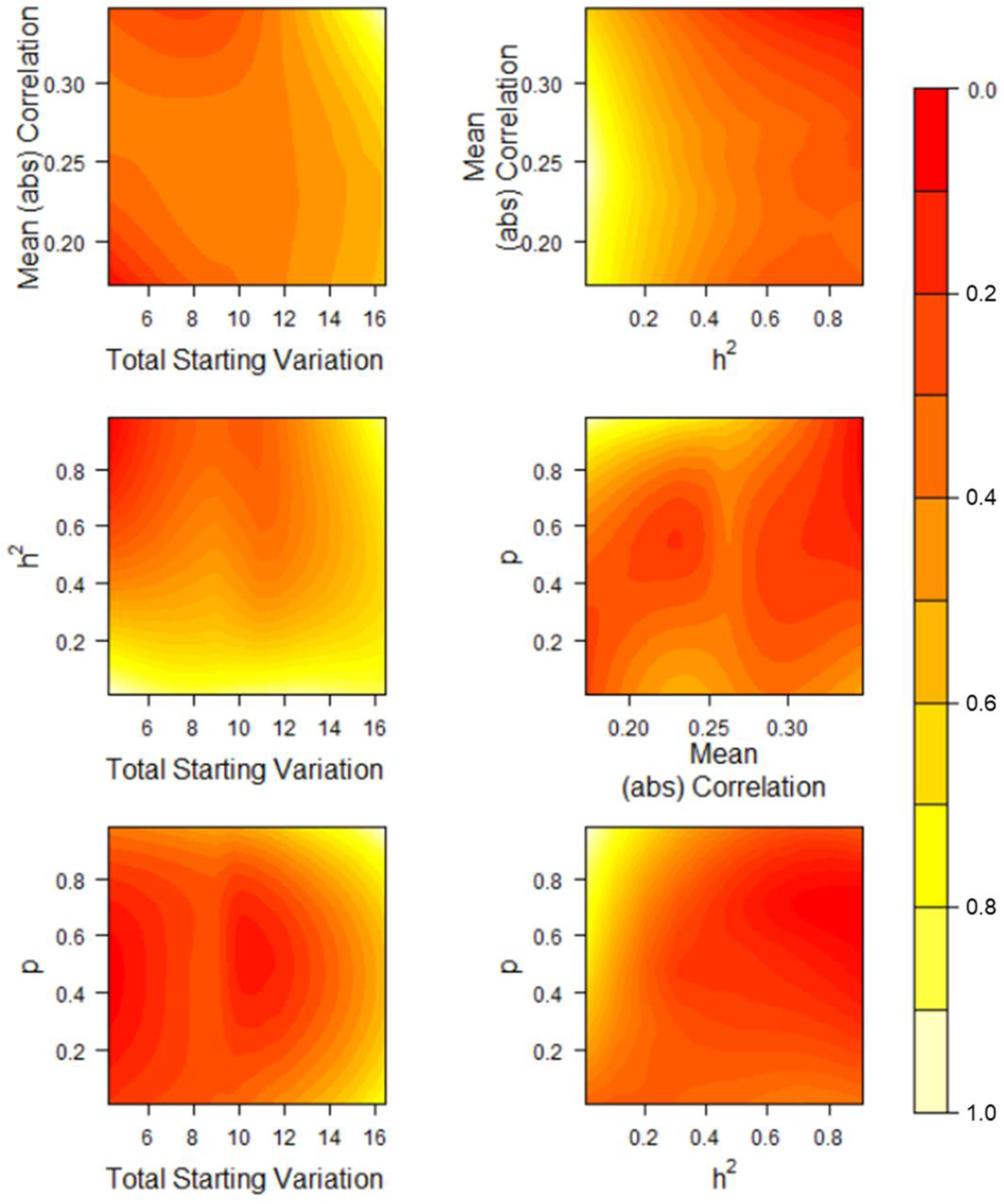
*λ*_2_⁄*λ*_1_ after evolution on holey landscapes remained low regardless of starting parameters (Table S10).

**Figure S12.**
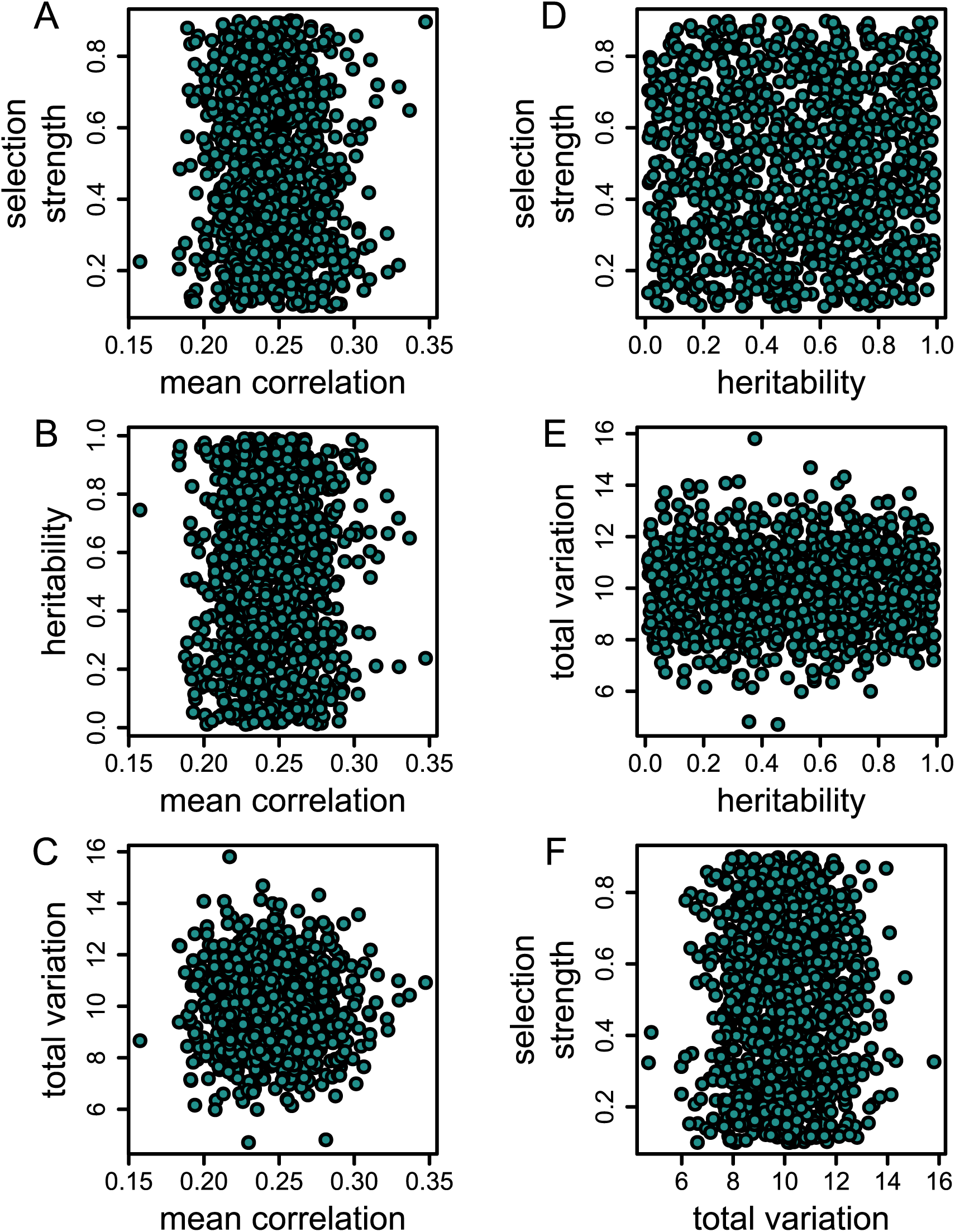
The parameter space across 1000 random sets of Gaussian landscapes covered a wide-range of plausible values of genetic variation, genetic correlations, and selection strengths. “Mean correlation” is the average *absolute* correlation (mean|*r*|) of an iteration’s 45 total correlations. The distribution of mean|*r*| is constrained due to values needing to satisfy the requirements to be a correlation matrix (e.g. being positive definite).

**Figure S13.**
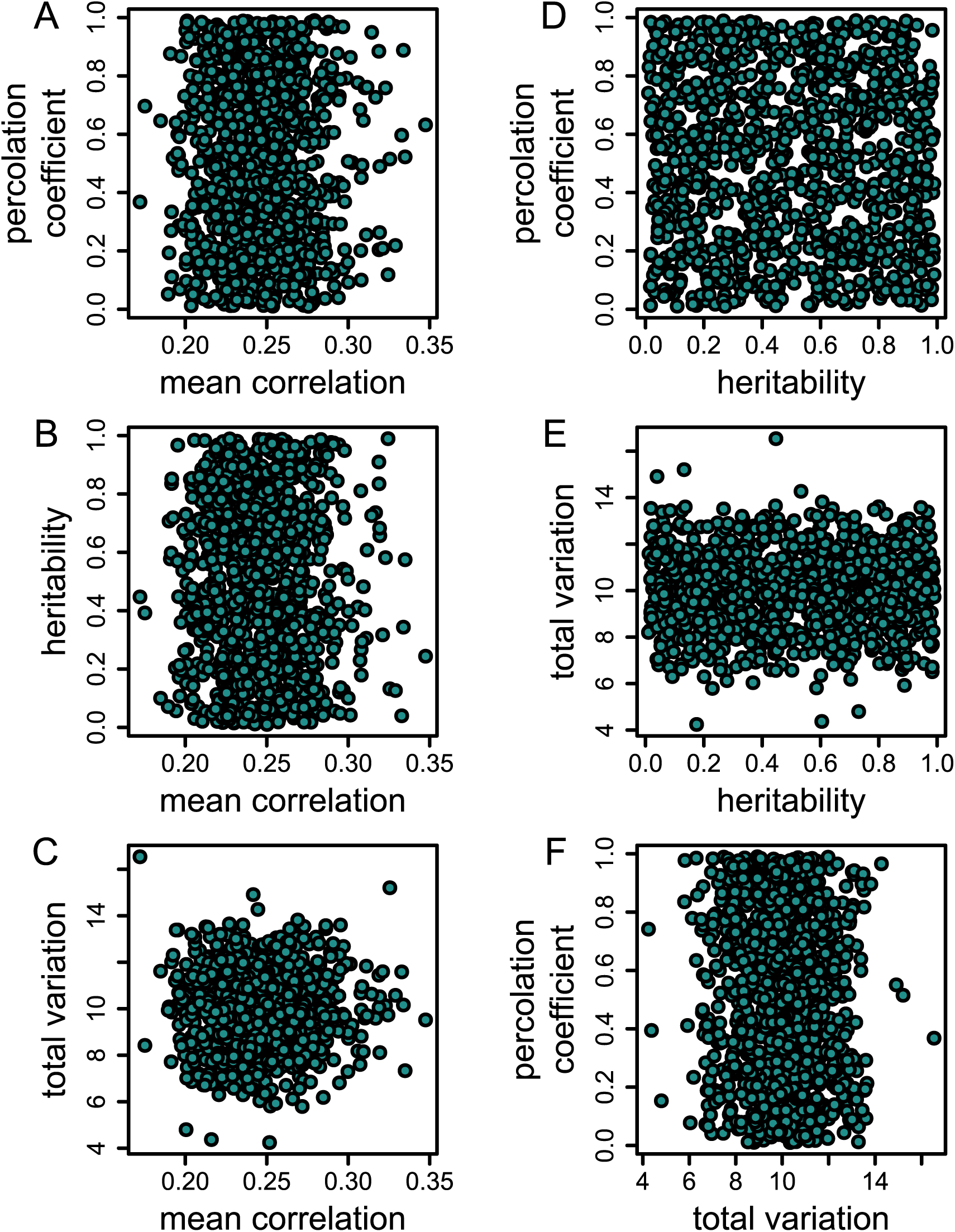
The parameter space across 1000 random sets of holey landscapes covered a wide-range of plausible values of genetic variation, genetic correlations, and proportion of viable phenotypes (i.e. the percolation coefficient). “Mean correlation” is the average *absolute* correlation (mean|*r*|) of the iteration’s 45 total correlations. The distribution of mean|*r*| is constrained due to values needing to satisfy the requirements to be a correlation matrix (e.g. being positive definite). The percolation coefficient (i.e. *p*) determines the probability that a phenotypic combination is viable.

**Figure S14.**
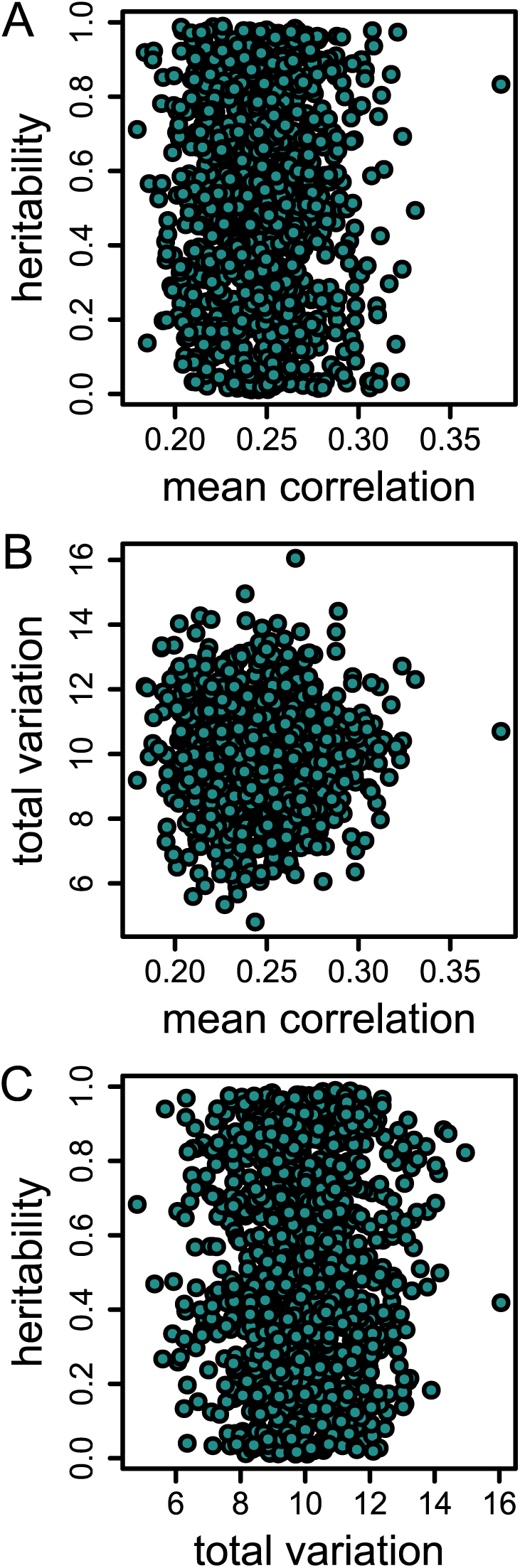
The parameter space across 1000 random sets of flat, drift landscapes covered a wide-range of plausible values of genetic variation and genetic correlations. “Mean correlation” is the average *absolute* correlation (mean|*r*|) of the iteration’s 45 total correlations. The distribution of mean|*r*| is constrained due to values needing to satisfy the requirements to be a correlation matrix (e.g. being positive definite).

**Figure S15.**
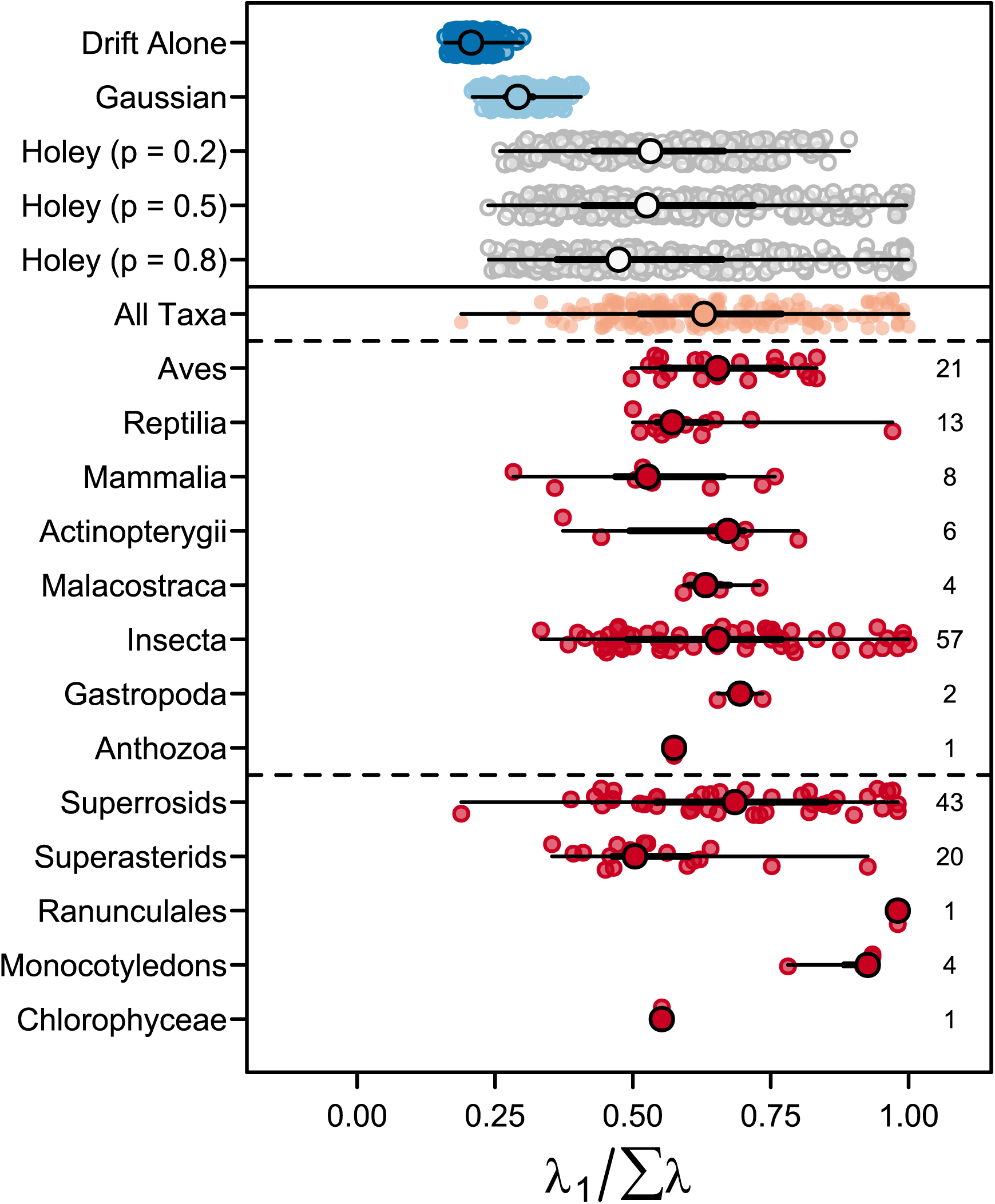
Modified “Orchard plot” of λ1/Σλ values for simulated (above solid line) and observed **G** matrices. *Trunks* (large points) are the medians for the specified group (e.g. Gaussian landscapes or Insecta), *branches* (thick lines) are interquartile ranges, *twigs* (thin lines) give the full range of values, and *fruits* (smaller points) are individual estimates within a simulation or taxonomic group. Rightmost numbers are the number of estimates available via literature search.

